# Geometric tortuosity at invaginating rod synapses slows glutamate diffusion and shapes synaptic responses: insights from anatomically realistic Monte Carlo simulations

**DOI:** 10.1101/2024.10.30.621088

**Authors:** Wallace B. Thoreson, Thomas M. Bartol, Nicholas H. Conoan, Jeffrey S. Diamond

## Abstract

At the first synapse in the vertebrate retina, rod photoreceptor terminals form deep invaginations occupied by multiple second-order rod bipolar and horizontal cell (RBP and HC) dendrites. Synaptic vesicles are released into this invagination at multiple sites beneath an elongated presynaptic ribbon. We investigated the impact of this complex architecture on the diffusion of synaptic glutamate and activity of postsynaptic receptors. We obtained serial electron micrographs of mouse retina and reconstructed four rod terminals along with their postsynaptic RBP and HC dendrites. We incorporated these structures into an anatomically realistic Monte Carlo simulation of neurotransmitter diffusion and receptor activation. We compared passive diffusion of glutamate in these realistic structures to existing, geometrically simplified models of the synapse and found that glutamate exits anatomically realistic synapses ten times more slowly than previously predicted. By comparing simulations with electrophysiological recordings, we modeled synaptic activation of EAAT5 glutamate transporters in rods, AMPA receptors on HC dendrites, and metabotropic glutamate receptors (mGluR6) on RRBP dendrites. Our simulations suggested that ~3,000 EAAT5 transporters populate the rod presynaptic membrane and that, while uptake by surrounding glial Müller cells retrieves much of the glutamate released by rods, binding and uptake by EAAT5 influences RBP response kinetics. The long lifetime of glutamate within the cleft allows mGluR6 on RBP dendrites to temporally integrate the steady stream of vesicles released at this synapse in darkness. Glutamate’s tortuous diffusional path through realistic synaptic geometry confers quantal variability, as release from nearby ribbon sites exerts larger effects on RBP and HC receptors than release from more distant sites. While greater integration may allow slower sustained release rates, added quantal variability complicates the challenging task of detecting brief decreases in release produced by rod light responses at scotopic threshold.

## Introduction

Rod photoreceptor synaptic terminals, termed spherules due to their bulbous shape, are structurally distinct from most other central synapses. The presynaptic active zone, which contains a plate-like ribbon structure tethering dozens of synaptic vesicles, apposes postsynaptic dendrites of multiple second-order neurons that terminate deep within an invagination into the rod spherule (Fig. 1A). Each synapse is typically occupied by two RBP dendrites and two HC dendrites that extend further into the invagination to flank the synaptic ridge. Absorption of even just a single photon by rhodopsin causes rods to hyperpolarize, temporarily decreasing the rate at which glutamate-filled vesicles are released at sites along the base of the ribbon (Moser et al., 2020; Thoreson, 2021). The resulting decrease in glutamate levels in the synaptic cleft alters the activity of glutamate receptors on RBP and HC dendrites. Here, we investigated how the complex architecture of this invaginating synapse influences the dynamics of glutamate diffusion following its release and, consequently, the synaptic responses of postsynaptic neurons.

**Fig. 1.**
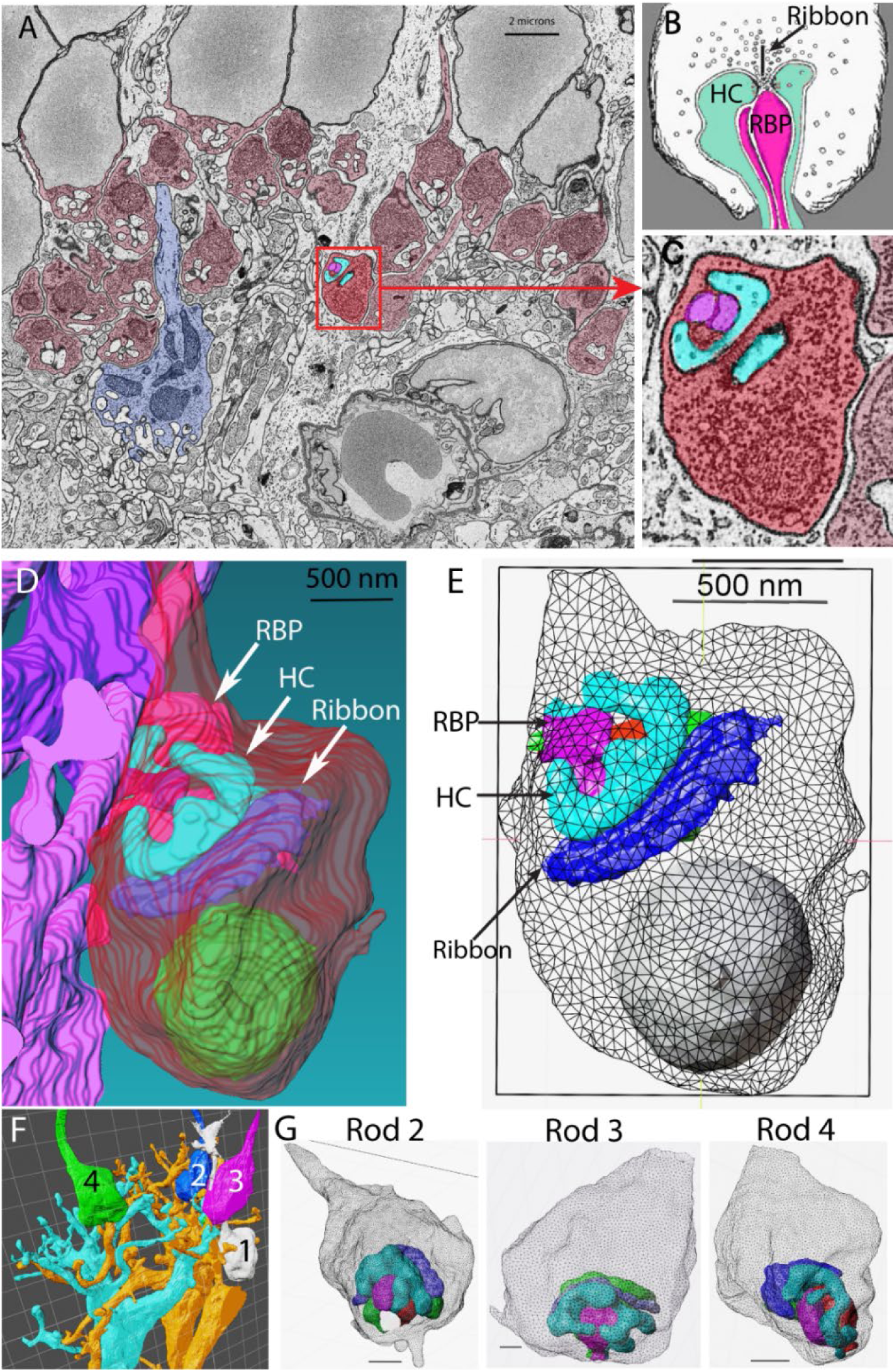
Model construction. A. Example serial block face scanning electron micrograph. Rod terminals colored in red; cone terminal colored in blue. Rod 1 used for reconstruction is shown in brighter red. B. Diagram of an invaginating rod synapse. Vesicles surround the synaptic ribbon. Rod bipolar cell (RBP) dendrites are magenta and HC dendrites are turquoise. Modified under a Creative Commons License from Webvision (http://webvision.med.utah.edu/) C. Magnified image of rod 1. Post-synaptic HC dendrites are turquoise and RBP dendrites magenta. D. Reconstructed rod 1 terminal along with HC and RBP dendrites. The synaptic ribbon (dark blue) and mitochondrion (green) are also shown. E. Mesh structures of the same cells. F. Illustration of four reconstructed rod spherules (rods 1-4) with two RBPs (blue and yellow). G. Mesh structures of rods 2-4 along with the post-synaptic HC and RBP dendrites. Details of these mesh structures are provided in Table 1.

**Table 1.** Details of mesh structures of rods, HC dendrites, and RBP dendrites used for simulations.

|  | <u>Rod 1</u> | <u>Rod 2</u> | <u>Rod 3</u> | <u>Rod 4</u> |
| --- | --- | --- | --- | --- |
| Surface Area ($\mu\text{m}^2$) | 29.15 | 27.67 | 32.29 | 26.13 |
| Volume ($\mu\text{m}^3$) | 5.94 | 5.024 | 6.576 | 4.771 |
| Faces | 6,580 | 14,498 | 57,802 | 30,384 |
| Cleft Volume ($\mu\text{m}^3$) | 0.564 | 0.572 | 0.6056 | 0.448 |
| Extracellular cleft volume ($\mu\text{m}^3$) | 0.1125 | 0.06 | 0.074 | 0.057 |
| Extracellular volume fraction | 0.2 | 0.11 | 0.12 | 0.13 |
| <u>Horizontal cell dendrite 1</u> |  |  |  |  |
| Surface Area ($\mu\text{m}^2$) | 2.758 | 3.272 | 2.726 | 2.382 |
| Volume ($\mu\text{m}^3$) | 0.1674 | 0.1945 | 0.1763 | 0.114 |
| Faces | 2040 | 4754 | 2388 | 1914 |
| <u>Horizontal cell dendrite 2</u> |  |  |  |  |
| Surface Area ($\mu\text{m}^2$) | 2.827 | 3.123 | 3.213 | 2.272 |
| Volume ($\mu\text{m}^3$) | 0.1894 | 0.1758 | 0.2228 | 0.1337 |
| Faces | 1088 | 7020 | 5530 | 2418 |
| <u>Bipolar cell dendrite 1</u> |  |  |  |  |
| Surface Area ($\mu\text{m}^2$) | 0.7964 | 1.2167 | 1.0288 | 1.1962 |
| Volume ($\mu\text{m}^3$) | 0.03475 | 0.07234 | 0.06593 | 0.07867 |
| Faces | 662 | 3084 | 5038 | 602 |
| <u>Bipolar cell dendrite 2</u> |  |  |  |  |
| Surface Area ($\mu\text{m}^2$) | 0.6539 | 1.041 | 1.0622 | 1.17 |
| Volume ($\mu\text{m}^3$) | 0.03208 | 0.069 | 0.06686 | 0.06456 |
| Faces | 1422 | 714 | 4832 | 568 |
| <u>Bipolar cell dendrite 2b</u> |  |  |  |  |
| Surface Area ($\mu\text{m}^2$) | 0.6792 | | | |
| Volume ($\mu\text{m}^3$) | 0.0276 | | | |
| Faces | 1732 |  |  |  |

Our understanding of glutamate dynamics at rod spherules has been shaped by work in which the dimensions of the rod synapse were measured and incorporated into geometrically simplified analytical diffusion models (Rao-Mirotznik et al., 1998; Rao-Mirotznik et al., 1995). The resulting analysis argued that glutamate is cleared from the synaptic cleft in only a few milliseconds, suggesting that vesicles act independently on post-synaptic dendrites as discrete quanta rather than being integrated over time in the cleft. It also suggested that postsynaptic RBP and HC dendrites all experience a similar rapid change in glutamate concentration following release of a synaptic vesicle by the rod. Here, we have tested whether these results hold in a diffusion model incorporating realistic synaptic architecture. We obtained block face scanning electron micrographs (SBFSEM) of mouse retina and reconstructed four rod spherules along with their postsynaptic contacts. We imported these realistic structures into MCell4, a Monte Carlo simulation program that represents neurotransmitter diffusion and receptor/transporter kinetics as stochastic processes (Franks et al., 2002; Garcia et al., 2022; Husar, 2022; Kerr et al., 2008; Stiles et al., 1996). We used MCell4 to evaluate effects of synaptic geometry on passive glutamate diffusion, glutamate uptake by transporters, and the activation of postsynaptic glutamate receptors.

Simulated rod membranes were populated with type 5 excitatory amino acid transporters (EAAT5) (Arriza et al., 1997; Eliasof et al., 1998; Gehlen et al., 2021; Pow and Barnett, 2000). EAAT5 retrieves a fraction of synaptically released glutamate (Hasegawa et al., 2006), although most uptake appears to be accomplished by EAAT1 in Müller glial membranes (Barnett and Pow, 2000; Derouiche, 1996; Eliasof et al., 1998; Fyk-Kolodziej et al., 2004; Harada et al., 1998; Pow et al., 2000; Rauen et al., 1996; Rauen et al., 1998; Sarthy et al., 2005). The EAAT5 transport cycle activates a large anion current (I_A(glu)_) (Arriza et al., 1997; Grant and Werblin, 1996; Schneider et al., 2014) that can be used as a reporter for presynaptic glutamate release (Hays et al., 2020; Szmajda and Devries, 2011). An existing kinetic model of EAAT2 (Kolen et al., 2020) was modified to represent EAAT5 and reproduce I_A(glu)_ recorded from rods in response to single vesicle release events (Kolen et al., 2020). Simulated HC membranes were populated with AMPA receptors (Bartol et al., 2015; Jonas et al., 1993; Lu et al., 2009), modified to reproduce miniature excitatory post-synaptic currents recorded in HCs (mEPSCs). RBP membranes were populated with mGluR6, which closes TRPM1 cation channels via a G_o_-type G protein-mediated intracellular signaling pathway (Koike et al., 2010a; Schneider et al., 2015), so that a reduction in glutamate release depolarizes RBPs.

Our simulations revealed that glutamate takes ten times longer to exit an anatomically realistic synaptic cleft than suggested previously by a geometrically simplified model (Rao-Mirotznik et al., 1998). This longer lifetime at postsynaptic receptors enhances temporal integration in the RBP and suggests that slower rod release rates than previously predicted can maintain the RBP membrane potential in darkness. Slower diffusion kinetics also leads to greater variability in glutamate concentration transients and receptor activation that depends strongly on the relative locations of the released vesicles along the ribbon and the post-synaptic dendrites in the cleft. These additional sources of variability complicate the challenging task facing RBPs of discriminating small rod light responses from synaptic noise, inviting a reconsideration of optimal strategies for detection at scotopic threshold.

## Methods

### Anatomical reconstructions and MCell simulations

Serial block face images were obtained using an Apreo VS scanning electron microscope (SEM; Thermo Scientific). The tissue was fixed by immersion with standard EM fixative (2% Glutaraldehyde, 2% Paraformaldehyde, 0.1M Phosphate Buffer). We sectioned a volume of 14.3 × 20.6 × 24 μm. Voxel dimensions are 6 × 6 × 40 nm and each image consisted of 3435 × 4000 pixels.

Three dimensional reconstructions were performed using Amira software. Reconstructed cell membranes were converted to 3-D triangular mesh structures using the CellBlender add-on to the 3D computer graphics program Blender (Fig. 1). The volume, surface area, and faces of the mesh structures for each of the 4 rods and their post-synaptic partners are provided in Table 1. Simulations were performed with these reconstructions using Monte Carlo simulation software (MCell) (Husar et al., 2024) that simulates molecular diffusion as a pseudo-random walk according to an assigned diffusion constant. Glutamate molecules diffuse in three dimensions in extracellular space and can interact stochastically with immobile receptors and transporters located on membrane surfaces. The behavior of each individual glutamate transporter and receptor was represented by stochastic transitions between states within an individual Monte Carlo model.

To simulate EAAT5 glutamate transporters, we adjusted the kinetics, density, and location of simulated transporters to reproduce the amplitude and time course of I_A(glu)_ recorded in rods in response to single vesicle release events. Simulated rod membranes were densely populated with EAAT5 (10,000/μm^2^ (Hasegawa et al., 2006) and represented by a simplified model for EAAT2 (Kolen et al., 2020). Glutamate binding and unbinding rates were adjusted to match the rising phase of averaged single-vesicle I_A(glu)_ events (Fig. S1A) and maintain the 10 μM EC_50_ measured for EAAT5 in mouse rods (Thoreson and Chhunchha, 2023). We tested ON-rates from 1×10^6^ M/s to 1×10^9^ M/s. ON-rates of 1×10^8^ M/s and 1×10^9^ M/s produced reasonable fits to the initial inward deflection in I_A(glu)_ (Fig. S1A). However, simulations with an ON rate of 1×10^9^ M/s were more sensitive to EAAT5 placement than fits with 1×10^8^ M/s (compare panels D and E in Fig. S2) and yielded fewer channel openings than predicted from single vesicle currents. To match the single-vesicle I_A(glu)_ decay when using an ON-rate of 1×10^8^, we increased by threefold the transition rate constant of T_out_*Glu → T_int_*Glu.

We examined effects of EAAT5 location on I_A(glu)_ kinetics by placing EAAT5 in four different regions: rod membrane deep within the invagination adjacent to RBP tips, the neck of invagination (Neck), surrounding the exit of the invagination (Exit), and distributed throughout the entire cleft (Cleft) (Fig. S1C). With an ON rate of 1×10^8^ M/s, placing EAAT5 adjacent to RBP dendritic tips (turquoise) or distributing transporters throughout the cleft (purple) both produced similar responses that provided a reasonable match to actual I_A(glu)_ events. Placing EAAT5 only in the neck of the invagination (green trace) or outside the cleft (deep blue) yielded decay kinetics that were faster than actual I_A(glu)_ events, as well as yielding fewer channel openings. Since distributing receptors throughout the cleft yielded comparable results to placing them deep within the invagination, we chose the former for subsequent simulations.

To estimate the number of EAAT5 transporters in rods, we began with evidence that single vesicle I_A(glu)_ events average 4.5 pA (Thoreson and Chhunchha, 2023). We calculated the anion driving force by plotting voltage-dependent changes in the average amplitude of single vesicle I_A(glu)_ events, fitting these data with a straight line. The reversal potential predicted from these measurements averaged −8.2 ± 0.62 mV (n=6 rods). The holding potential of −70 mV therefore generates a driving force of 62 mV. The single channel conductance of EAAT5 measured with nitrate as a charge carrier was found to be 0.6 pS (Schneider et al., 2014) and glutamate transporters in salamander rods show a single channel conductance with chloride as the charge carrier of 0.7 pS (Larsson et al., 1996). SCN^−^ is more permeable than chloride or nitrate, so we assumed a larger single channel conductance of ~1 pS. These values suggest that ~72 anion channels are open at the peak of a single vesicle release event.

The maximum open probability of EAAT2 anion channels has been estimated at ~0.06 (Kolen et al., 2020). To calculate the open probability for anion channels achieved in our simulations, we divided the number of open anion channels by the total number of glutamate-bound states. We used the same rate for entry into the open state from T_int_*Glu as in the earlier model for EAAT2 (9,566/s) (Kolen et al., 2020). Assuming that channels are equally likely to exit the open state (i.e., open state ◊ T_int_*Glu = 9566/s), we obtain an open probability of 0.077. This seems reasonable given that EAAT5 anion currents are thought to have a larger open probability than EAAT2.

Like other metabotropic glutamate receptors, mGluR6 forms a homodimer and both members need to bind glutamate for G-protein activation (Levitz et al., 2016; Pin and Acher, 2002). We therefore modeled mGluR6 activation as two sequential glutamate-binding steps, considering the doubly bound mGluR6 dimer to be the activated receptor (Fig. S2). We assumed that the decay in the number of active mGluR6 receptors (i.e., decay of the doubly bound state) following release of a single vesicle should be at least as fast as the inward current evoked in RBPs by a saturating light flash (20-80% increase: 25.4 ± 1.9 ms; SEM, n=19; J. Pahlberg, unpublished). As illustrated in Fig. S2, we tested different rate constants and achieved a similar rate of decay in the doubly bound state using an OFF rate for glutamate unbinding of 500/s (20-80%: rod 1, 27 ms; rod 2, 50 ms; rod 3, 36 ms; rod 4, 28 ms). Combining this OFF rate with an ON rate for the initial glutamate binding step of 1e^8^ M/s yielded a steady state EC_50_ of 14 μM (Fig. S2), matching that measured by the displacement of glutamate from mGluR6 (12.3 μM; (Pin and Acher, 2002)). These parameters also yielded a slope factor of 1.4, similar to changes in mGluR6-mediated responses as a function of light intensity in dark-adapted RBPs (Berntson et al., 2004; Sampath and Rieke, 2004).

To simulate HC AMPA receptors, we used a kinetic model of AMPARs empirically derived from hippocampal neurons (Jonas et al., 1993). The original parameters were developed to fit data obtained at room temperature and later modified for a temperature of 34°C (Bartol et al., 2015).

Release from photoreceptors varies linearly with I_Ca_ (Thoreson et al., 2004) and so we estimated voltage-dependence of release rates in rods from changes in I_Ca_. For the voltage-dependence of I_Ca_, we used a Boltzmann function modified for driving force with an activation midpoint (V_50_) obtained from a sample of mouse rods (−31.5 mV; n=8) along with a reversal potential of +41 mV (Grassmeyer and Thoreson, 2017) and slope factor of 9 (Haeseleer et al., 2016).

Other analysis and data visualization procedures were performed using Clampfit 10, GraphPad Prism 9, Adobe Illustrator, and Adobe Photoshop software.

### Mice

For electrophysiology studies, we used mice of both sexes aged between 4-8 weeks. Euthanasia was performed by CO_2_ asphyxiation followed by cervical dislocation in accordance with AVMA Guidelines for the Euthanasia of Animals. Animal care and handling protocols were approved by the University of Nebraska Medical Center Institutional Animal Care and Use Committee. Experiments were conducted using C57BL/6J mice.

### Whole cell recordings

Whole cell recordings of rods were obtained using flatmount preparations of isolated retina. Eyes were enucleated after euthanizing the mouse and placed in Ames’ medium (US Biological; RRID:SCR_013653) bubbled with 95% O2/5% CO2. The cornea was punctured with a scalpel and the anterior segment removed. The retina was isolated after cutting optic nerve attachments. We then made three or four fine cuts at opposite poles and flattened the retina onto a glass slide in the perfusion chamber with photoreceptors facing up. The retina was anchored in place with a brain slice harp (Warner Instruments, cat. no. 64-0250). To expose rod inner segments in flatmount retina, we gently touched the photoreceptors with a piece of nitrocellulose filter paper and then removed it to pull away adherent outer segments. The perfusion chamber was placed on an upright fixed-stage microscope (Nikon E600FN) equipped with a 60x water-immersion, long-working distance objective (1.0 NA). The tissue was superfused with room temperature Ames’ solution bubbled with 95% O_2_/5% CO_2_ at ~1 mL/min.

Whole cell recordings were performed using either an Axopatch 200B amplifier (Molecular Devices) with signals digitized by a DigiData 1550 interface (Molecular Devices) using PClamp 10 software or Heka EPC-10 amplifier and Patchmaster software (Lambrecht, Pfalz, Germany). Currents were acquired with filtering at 3 kHz.

Patch recording electrodes were pulled on a Narishige (Amityville, NY) PP-830 vertical puller using borosilicate glass pipettes (1.2 mm outer diameter, 0.9 mm inner diameter with internal filament; World Precision Instruments, Sarasota, FL). Pipettes had tip diameters of 1–2 μm and resistances of 10–15 MΩ. Rod inner segments were targeted with positive pressure using recording electrodes mounted on Huxley-Wall or motorized micromanipulators (Sutter Instruments, MP225).

Rod ribbons are surrounded by the glutamate transporter EAAT5 (Arriza et al. 1997; Eliasof et al. 1998; Hasegawa et al. 2006) and glutamate reuptake into rods by these transporters activates a large, anion current (I_A(glu)_) (Arriza et al., 1997; Grant and Werblin, 1996; Schneider et al., 2014). I_A(glu)_ is thermodynamically uncoupled from the transport process (Machtens et al., 2015). Glutamate transporter anion currents can be observed in rods using Cl^−^ as the principal anion (Hays et al., 2020), but are enhanced by replacing Cl^−^ with a more permeable anion like thiocyanate (SCN^−^) in the patch pipette (Eliasof and Jahr, 1996). The intracellular pipette solution for these experiments contained (in mM): 120 KSCN, 10 TEA-Cl, 10 HEPES, 1 CaCl2, 1 MgCl2, 0.5 Na-GTP, 5 Mg-ATP, 5 EGTA, 5 phospho-creatine, pH 7.3. Voltages were not corrected for a liquid junction potential of 3.9 mV.

To record mEPSCs from HCs, horizontal slices of retina were prepared as described elsewhere (Feigenspan and Babai, 2017). Briefly, retinas were isolated and then embedded in 1.8% low gelling agarose (Sigma-Aldrich). Horizontal slices (200 μm thick) were cut parallel to the plane of the retina using a vibratome (Leica Microsystems) at room temperature.

For HC recordings, we used a pipette solution containing (in mM): 120 KGluconate, 10 TEACl, 10 HEPES, 5 EGTA, 1 CaCl2, 1 MgCl2, 0.5 NaGTP, 5 MgATP, 5 phosphocreatine, pH 7.2-7.3. HCs were identified visually and confirmed physiologically by the characteristic voltage-dependent currents, particularly prominent A-type K^+^ currents (Feigenspan and Babai, 2017). In our initial recordings, HC identity was confirmed anatomically by loading the cell with the fluorescent dye Alexa 488 (Invitrogen, Waltham, MA) through the patch pipette. Chemical reagents were obtained from Sigma-Aldrich unless otherwise indicated.

## Results

### Model construction

To create an anatomically detailed diffusion model of rod photoreceptor synapses, we first obtained a series of serial block face scanning electron microscope (SBSFSEM) images from the outer retina of a C57Bl6J mouse (Fig. 1A, B). By viewing consecutive sections at high magnification, we reconstructed four rod spherules along with their postsynaptic HC and RBP dendrites (Fig. 1C). We then rendered the reconstructed synapses as a collection of 3D surfaces (Fig. 1D) and imported them into MCell. The four reconstructed terminals had the same general structure but exhibited significant variability in geometric dimensions (Fig. 1E-G; Table 1).

Rao-Mirotznik et al. (1998) modeled the cat rod synapse as a sphere with a narrow neck for the exit. We created similar models in MCell, configuring spheres with volumes to match the clefts of mouse rods 1 and 2 (Fig. 2A, sphere models shown at the same scale as reconstructed terminals). Like the earlier model, these spheres emptied through a narrow neck (r= 0.12 μm, length = 0.1 μm).

**Fig. 2.**
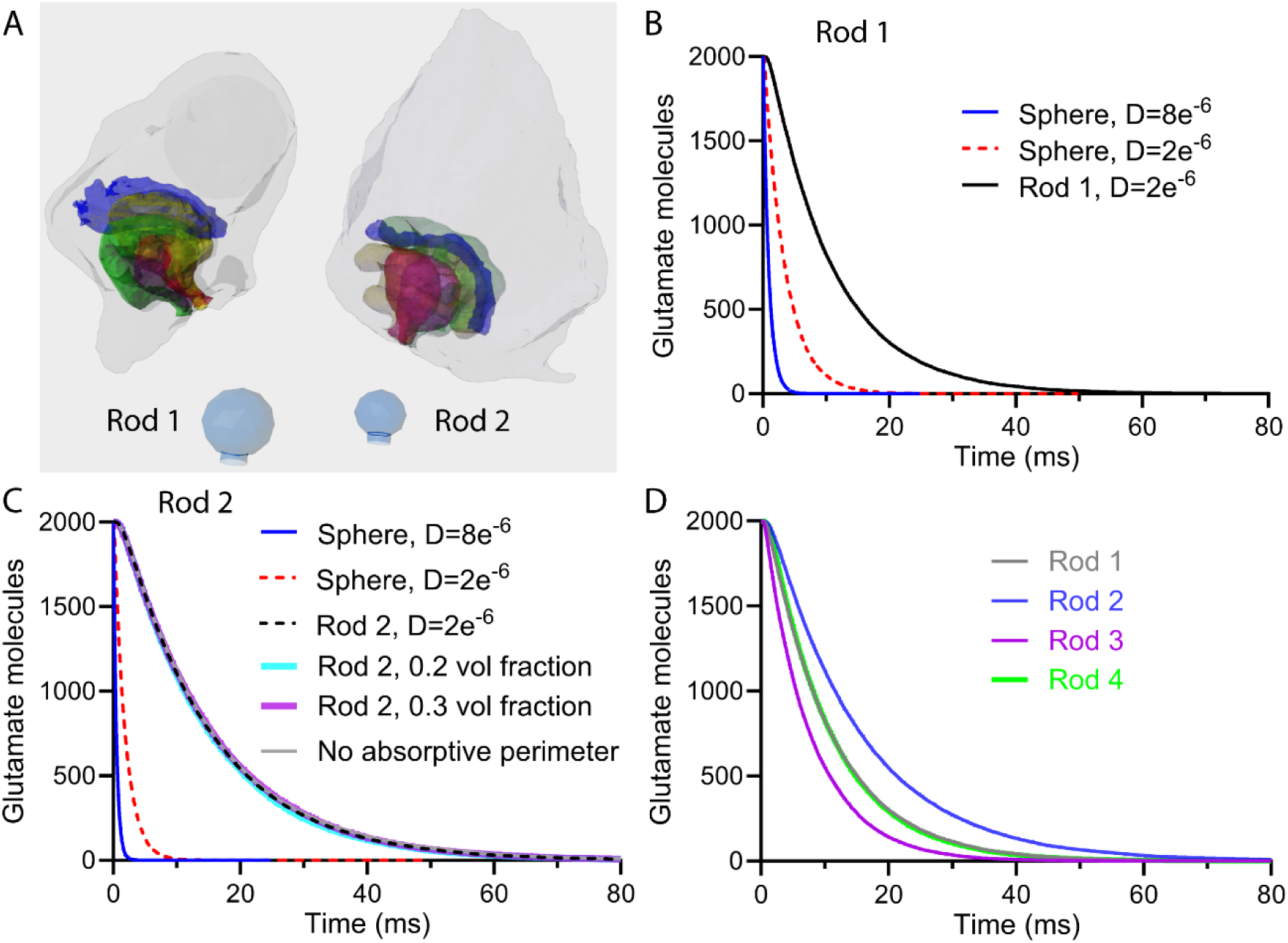
Simulations of passive glutamate diffusion comparing reconstructed rod spherules with simplified models that represented the extracellular synaptic volume as a sphere with a narrow neck for an exit. A. Illustration of reconstructed rod spherules 1 and 2. HC dendrites are yellow and green, bipolar cell dendrites are red and magenta, and ribbons are blue. Below each spherule is the corresponding sphere model containing the same extracellular volume shown at the same scale. B. Plot of the number of glutamate molecules that remained after release of a vesicle in the synaptic cleft of rod 1 (D = 2 × 10^−6^ cm^2^/s, black line) or a sphere with the same extracellular volume (D = 8 × 10^−6^ cm^2^/s, blue line; D = 2 × 10^−6^ cm^2^/s, dashed red line). We simulated release in rod 1 at a site just beneath the center of the ribbon. C. Graph of the glutamate decline in a sphere with the same extracellular volume as the invaginating synapse of rod 2 (D = 8 × 10^−6^ cm^2^/s, blue line; D = 2 × 10^−6^ cm^2^/s, dashed red line). This graph also shows the decline in glutamate following release of a vesicle beneath the center of the ribbon in rod 2 with volume fractions of 0.11 (D = 2 × 10^−6^ cm^2^/s, black line), 0.2 (turquoise line), and 0.3 (purple line). Eliminating an absorptive perimeter that simulates avid Muller cell uptake had no effect on the time course of decay (gray trace). D. Comparison of passive glutamate decline in the four rod spherules. Fitting the decays with single exponentials yielded the following time constants: rod 1: 10.64 ms, rod 2: 14.10 ms, rod3: 7.45 ms, rod 10: 8.89 ms. Each trace is an average of 12 simulations run using different seed values.

Estimates of the number of glutamate molecules filling each synaptic vesicle vary widely (Orrego and Villanueva, 1993; Takamori et al., 2006; Wang et al., 2019), so we chose a moderate value of 2,000 glutamate molecules. As mouse rod synaptic vesicles have an inner diameter of about 30 nm (Fuchs et al., 2014), 2000 glutamate molecules would constitute approximately 250 mM, close to biochemical measurements of purified synaptic vesicles from cortex (210 mM; (Riveros et al., 1986).

### Effects of synaptic geometry on passive diffusion of glutamate

We first simulated the release of 2,000 glutamate molecules from points at the apexes of the two spheres containing the cleft volumes of rods 1 and 2. Using a diffusion coefficient (D) describing free diffusion of glutamate in saline (D = 8 × 10^−6^ cm^2^/s; Rao-Mirotznik et al., 1998), our Monte Carlo simulations predicted rapid clearance from the synaptic cleft (τ = 0.88 and 0.46 ms for spheres 1 and 2, respectively; Figs. 2B,C). Using a larger sphere that matched the cleft volume of a cat rod spherule (0.21 μm^3^; Rao-Mirotznik, et al., 1995), Monte Carlo simulations yielded a 1.6 ms time constant for glutamate clearance, close to the value of 1.7 ms predicted analytically by Rao-Mirotznik et al. (1998).

Work at other central synapses indicates that glutamate diffusion in extracellular space at synapses is slower than in free solution (Nicholson and Hrabetova, 2017; Nicholson et al., 1979; Nielsen et al., 2004; Rusakov and Kullmann, 1998; Sykova, 2004). Lowering D to account for the viscous tortuosity of extracellular space from 8 to 2 × 10^−6^ cm^2^/s (Franks et al., 2002) slowed diffusion and consequently glutamate clearance several fold (Figs. 2B, C).

We next analyzed the impact of realistic synaptic geometry by comparing results of the two sphere models to simulations of single vesicle release in the reconstructed rod synapses. Within the realistic geometry, we positioned the vesicle release site just beneath the ribbon at the center of the invagination. These simulations produced even slower glutamate clearance time constants (Fig. 2B,C). Notably, the cross-sectional area of extracellular space at the neck of the synaptic invaginations closely approximated that of the exit from the simplified spherical models, indicating that slower clearance from the reconstructed synapses was not due to greater constriction at the neck.

Given the larger extracellular volume fraction (α) of rod 1 compared to rod 2 (Table 1), we tested the effects of this parameter on our results. To do so, we shrank the post-synaptic dendrites in rod 2 to increase α from 0.11 to 0.2 and 0.3 and found that the time constant of glutamate decay remained the same (Fig. 2C). Taken together, these results show that the geometric tortuosity introduced by the complicated anatomy of the invaginating rod synapse dictates the dynamics of neurotransmitter diffusion (Nicholson and Hrabetova, 2017; Nielsen et al., 2004; Rusakov and Kullmann, 1998).

Müller glial cell processes envelope rod synapses and retrieve glutamate molecules that escape the cleft (Attwell et al., 1989; Rauen et al., 1998; Sarantis and Mobbs, 1992). To simulate avid Müller cell uptake, we wrapped the entire rod terminal with an absorptive mesh to remove any glutamate molecule that exited the synaptic cleft immediately. This absorptive perimeter had no effect on the time course of glutamate clearance compared to simulations in which escaping glutamate entered a large open volume (Fig. 2C). This suggests that, while Müller cell uptake is likely important for maintaining a steep glutamate diffusion gradient at the mouth of the synapse, it does not directly regulate the rate of glutamate clearance from the cleft.

We also examined passive glutamate diffusion at the other two reconstructed rod synapses. Rods 3 and 4 had similar cleft volumes and α compared to rod 2 (Table 1). As observed with rod 2, increasing α of rods 3 and 4 did not substantially affect the rate of glutamate clearance. Although rods 2, 3 and 4 all had similar cleft volumes, the rates of glutamate clearance ranged from 7.4 ms to 14 ms among them (Fig. 2D). Interestingly, rods 1 and 4 exhibited similar simulated glutamate clearance rates, even though rod 1 has twice the extracellular volume (Fig. 2D). These rod-to-rod differences further demonstrate the powerful influence of realistic geometric tortuosity on the kinetics of glutamate diffusion within invaginating rod synapses (Nicholson and Hrabetova, 2017; Rusakov and Kullmann, 1998). The combined effects of viscous and geometric tortuosity slow glutamate clearance from rod synapses tenfold compared to exit from a saline-filled sphere. The unexpectedly prolonged presence of glutamate in the cleft prompted us to simulate its interaction with synaptic glutamate transporters and receptors.

To predict glutamate concentrations achieved at RBP dendrites, Rao-Mirotznik et al. modeled the invaginating rod synapse as three slabs intersecting at 120 degrees, with a vesicle release site positioned at the confluence of the slabs. Fig. 3A shows a schematic of the invaginating synapse with a rod bipolar cell (RBP) dendrite terminating some distance below the ribbon release site while the two HC dendrites flank the synaptic ridge. The model simulates this structure by using two slabs to represent the extracellular space between the rod and HC membranes while the third slab represents the extracellular space above a RBP terminal (Fig. 3A). We recreated this arrangement in MCell with each slab consisting of two planes separated by 16 nm, then simulated release of 2000 glutamate molecules at the vertex of this narrow cleft. We measured the number of glutamate molecules that entered a small measurement box (15 × 100 × 200 nm) placed 70 nm from the release site (Fig. 3A). As predicted by Rao-Mirotznik et al., Monte Carlo simulations of release of 2,000 molecules showed an abrupt rise and rapid decline of glutamate within this region (Fig. 3C). We compared this model with the four reconstructed synapses by placing a measurement box (15 × 100 × 200 nm) just above RBP dendrites (e.g., Fig. 3B). The proximity of the measurement box to the release site minimized effects of tortuosity and so we saw a similarly rapid rise and fast decay in the rod synapses (τ_fast_ = 0.114 ± 0.0217 ms; τ_slow_ = 0.589 ± 0.1394 ms; Fig. 4C) and the planar model (τ_fast_ = 0.081 ms; τ_slow_ = 0.558 ms). We converted the number of molecules in the measurement regions to concentration (Fig. 3C). The peak concentration attained in the planar model was slightly lower than that attained in synapses (0.775 ± 0.3325 mM) but both are consistent with estimates of synaptic glutamate levels at rod synapses obtained by use of low affinity antagonists (Cadetti et al., 2008; Kim and Miller, 1993).

**Fig. 3.**
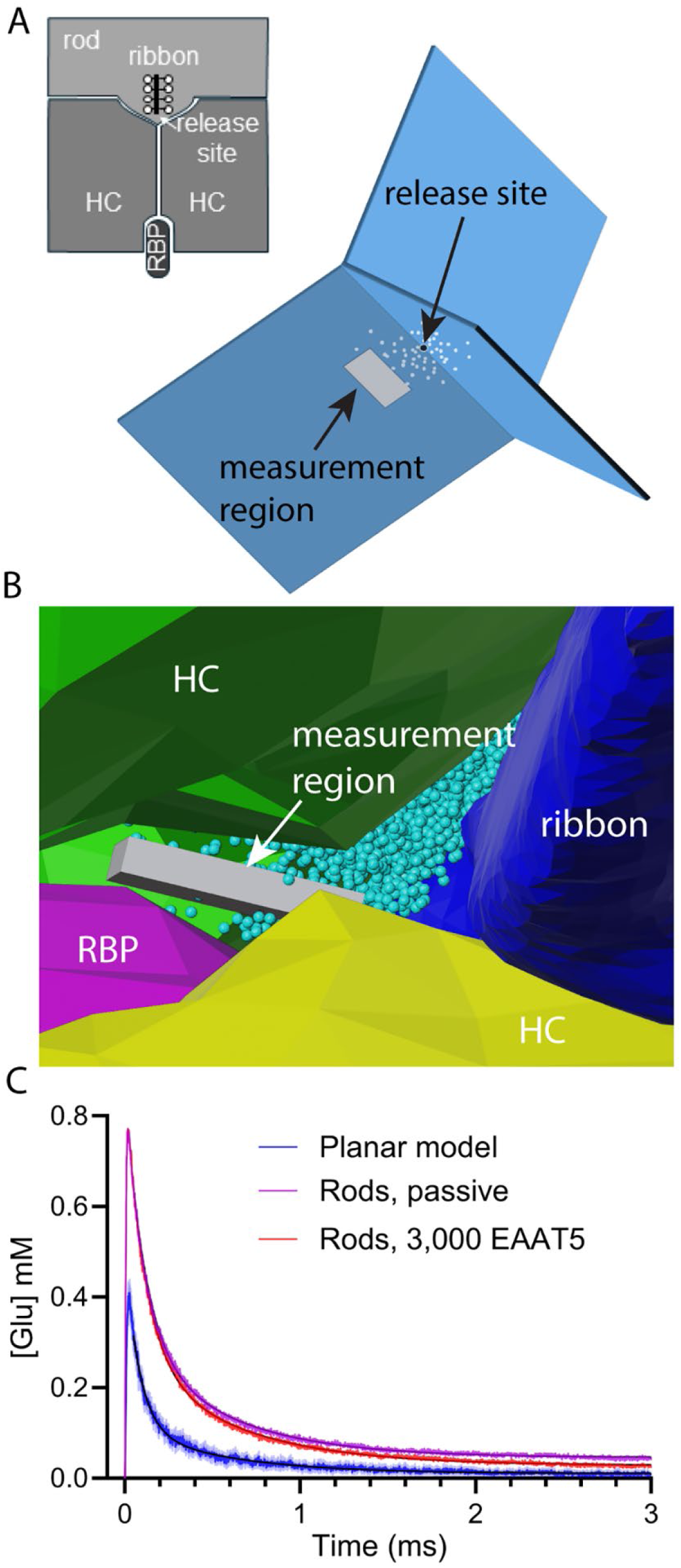
Comparing glutamate kinetics between a planar model and reconstructed rod spherules. A. Schematic representation of an invaginating rod synapse that shows two horizontal cell (HC) dendrites flanking the synaptic ridge (upper left) and a rod bipolar cell dendrite (RBP) terminating beneath the ribbon release site. The image at the lower right shows a simplified planar model of this synaptic arrangement that consists of three slabs intersecting at an angle of 120 degrees. Each slab involves two planes separated by 16 nm to simulate a synaptic cleft. We simulated release of 2000 glutamate molecules at the vertex of this narrow cleft. We measured the number of glutamate molecules that entered a small measurement region (gray box, 15 × 100 × 200 nm) with the leading edge 70 nm from the release site. B. Illustration of a reconstructed synapse (rod 1) with a measurement region (gray box, 15 × 100 × 200 nm) placed just above a RBP dendrite (purple). HC dendrites are shown in yellow and green. The ribbon is dark blue and glutamate molecules are sky blue. C. Monte Carlo simulations of single vesicle release showed an abrupt rise and rapid decline of glutamate in measurement regions in the planar model (blue trace; average of 12 seeds) and four reconstructed synapses (purple trace, average of four rods, 12 seeds apiece). Red trace shows the change in glutamate observed when the rod models included active uptake by 3,000 EAAT5 distributed throughout the synaptic cleft. The measurement regions attained a peak concentration of 0.4 mM in the planar model and an average of 0.775 ± 0.3325 mM in the four rod synapses. Decay kinetics were fit with two exponentials. Planar model: τ_fast_ = 0.081 ms; τ_slow_ = 0.558 ms. Passive glutamate decay in the four rod synapses: τ_fast_ = 0.114 ± 0.0217 ms; τ_slow_ = 0.589 ± 0.1394 ms. Decay in the rod synapses with active uptake by 3,000 EAAT5: τ_fast_ = 0.116 ± 0.0238 ms; τ_slow_ = 0.644 ± 0.1943 ms.

**Fig. 4.**
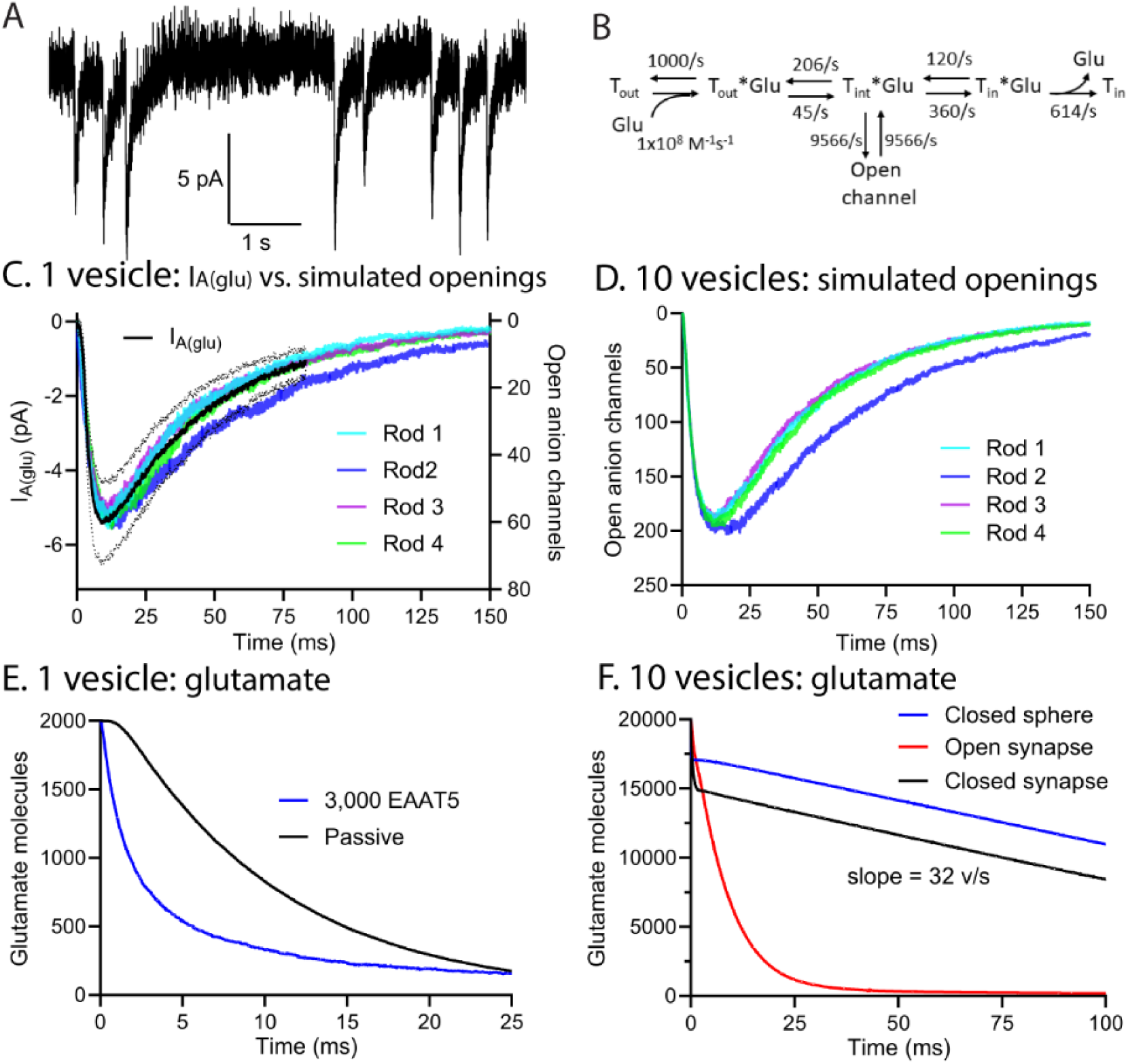
Simulations of EAAT5 anion channel activity. A. Example of EAAT5 anion currents evoked by single vesicle release events in a rod. B. Reaction scheme for EAAT5 (modified from a model for EAAT2)(Kolen et al., 2020). C. Colored traces show the average simulated EAAT5 anion channel activity in four rods following release of a single vesicle. Black trace shows the average change in I_A(glu)_ (± S.D.) evoked by single vesicle release events in 15 rods (7-72 events/rod). D. EAAT5 anion channel activity evoked in four rods by simulating simultaneous release of 10 vesicles. E. Comparison of the decline in glutamate molecules following simulated release of a single vesicle with (blue trace) and without (black trace) 3,000 EAAT5 transporters in rod 2. F. Graph of the decline in glutamate following simultaneous release of 10 vesicles in rod 2 with open (red trace) or closed (black trace) exits at the mouth of the synapse. The decline in a closed sphere with the same volume as rod 2 is plotted for comparison (blue trace). All simulations are the average of 12 runs with different seed values.

### Simulations of the glutamate transporter, EAAT5

We next examined the influence of EAAT5 glutamate transporters, the principal glutamate transporters in rods (Arriza et al., 1997; Eliasof et al., 1998) on glutamate lifetime in the synaptic cleft (Fig. 4). Modifying a model for EAAT2 (Kolen et al., 2020), we adjusted the kinetics, density, and location of simulated transporters to reproduce the amplitude and time course of I_A(glu)_ recorded in rods in response to single vesicle release events (Fig. 4; see Methods). The best fit to recorded EAAT5 currents was obtained by placing EAAT5 in the rod membrane within the synaptic cleft (see Methods), consistent with immunohistochemical localization of this protein (Gehlen et al., 2021).

Rod I_A(glu)_ responses evoked by photolytic uncaging of glutamate or strong depolarizing voltage steps reach a maximum of 12-13 pA (Mesnard et al., 2022b; Thoreson and Chhunchha, 2023), only three times larger than single vesicle events, suggesting that EAAT5 transporters can be saturated by simultaneous release of as few as 3 vesicles. With this constraint in mind, we simulated release of 10 vesicles and progressively reduced the number of EAAT5 transporters until we achieved a state where responses showed saturating responses equivalent to 3-4 vesicles. With 3,000 EAAT5 transporters distributed throughout the synaptic cleft, a single vesicle stimulated ~60 open anion channel openings while simultaneous release of 10 vesicles opened ~190 channels, showing saturation after release of slightly more than 3 vesicles (Fig. 4D). We therefore proceeded with simulations containing 3,000 EAAT5 transporters in the rod membrane.

Fig. 4E shows the impact on synaptic glutamate levels of uptake by 3,000 EAAT5 transporters in rod 2. Following release of a single vesicle, the simulations show that rapid binding of glutamate to EAAT5 speeds the initial decline in free glutamate. This is followed by a slower decline dictated by the rate of diffusion out of the synaptic cleft (τ_fast_ = 1.8 ms; τ_slow_ = 15 ms; n=12 trials). For a single vesicle release event, the fast component accounted for 73% of the total decline in glutamate, consistent with binding of 1,460 glutamate molecules by 3,000 transporters. Rods are capable of multivesicular release events consisting of 10 or more vesicles (Hays et al., 2021). The presence of glutamate uptake by EAAT5 had only small effects on the kinetics of glutamate reaching RBP dendrites. Including 3,000 EAAT5 slightly accelerated glutamate decay in the measurement region placed just above RBP dendrite in the previous figure (Fig. 4C; τ_fast_ = 0.12 ms; τ_slow_ = 0.56 ms). When we simulated simultaneous release of 10 vesicles, the fast component corresponding to glutamate binding of EAAT5 constituted a much smaller proportion of the total decline since the number of glutamate molecules (20,000) was much greater than the number of transporters (Fig. 4F)

To measure the maximum rate of glutamate uptake by 3,000 EAAT5 transporters, we simulated release of 10 vesicles inside a closed synaptic cleft (rod 2; Fig. 4F). The only exit available to glutamate in this simulation was uptake by EAAT5. Uptake settled to a constant rate of 63,390 glutamate molecules/s or 32 vesicles/s. Performing the same simulation in a closed sphere with the volume of rod 2 yielded the same uptake rate. While EAAT5 can retrieve glutamate at rates up to 32 vesicles/s, glutamate declines much more rapidly when the synapse remains open, indicating that most of the glutamate diffuses out of the synaptic cleft, to be retrieved by Müller glia. Thus, our evidence suggests that EAAT5 in rods can take up functionally significant amounts of glutamate, as proposed by (Hasegawa et al., 2006), but most of it is likely to be retrieved by extra-synaptic Müller cells, as suggested by others (Harada et al., 1998; Niklaus et al., 2017; Pow et al., 2000; Rauen et al., 1998; Sarthy et al., 2005).

### Simulations of mGluR6 receptors on RBPs

Glutamate released from rods acts at mGluR6 receptors on RBP dendrites (Nomura et al., 1994). Activating these receptors triggers a signaling cascade that leads to closing of TRPM1 cation channels (Koike et al., 2010b; Morgans et al., 2010). The signaling cascade is not understood in sufficient detail for a complete model so we limited our model to the binding of glutamate to mGluR6. Class 3 metabotropic glutamate receptors--including mGluR6—form obligate homodimers in which ligand binding to both members is needed to activate the G protein cascade (Levitz et al., 2016; Pin and Bettler, 2016). We placed 200 receptors at the tips of each of the two bipolar cell dendrites and modeled receptor activation as two sequential glutamate-binding steps, considering the doubly bound mGluR6 dimer to be the activated receptor (Fig. S2).

We compared simulations of mGluR6 activity in reconstructed rods to mGluR6 activity in the corresponding sphere model (Fig. 5). We placed 400 receptors in a transparent plane adjacent to a release site at the apex of the sphere matching the cleft volume of rod 2 (Fig. 5A). This sphere is also close to cleft volumes of rods 3 and 4 (Table 1). We tested mGluR6 binding with and without 3,000 EAAT5 transporters placed on the inner surface of the sphere. Nearly all of the mGluR6 receptors in the sphere were rapidly bound following the release of a single glutamate-filled vesicle, and mGluR6 activation then decayed with a single time constant (10.1 ms; Fig. 5C, Table 2) that actually became faster when EAAT5 was removed (6.6 ms; Fig. 5D, Table 2). The slower decay in the presence of EAAT5 suggests that the transporters buffer glutamate, delaying its escape from the cleft and prolonging its interaction with mGluR6.

**Fig. 5.**
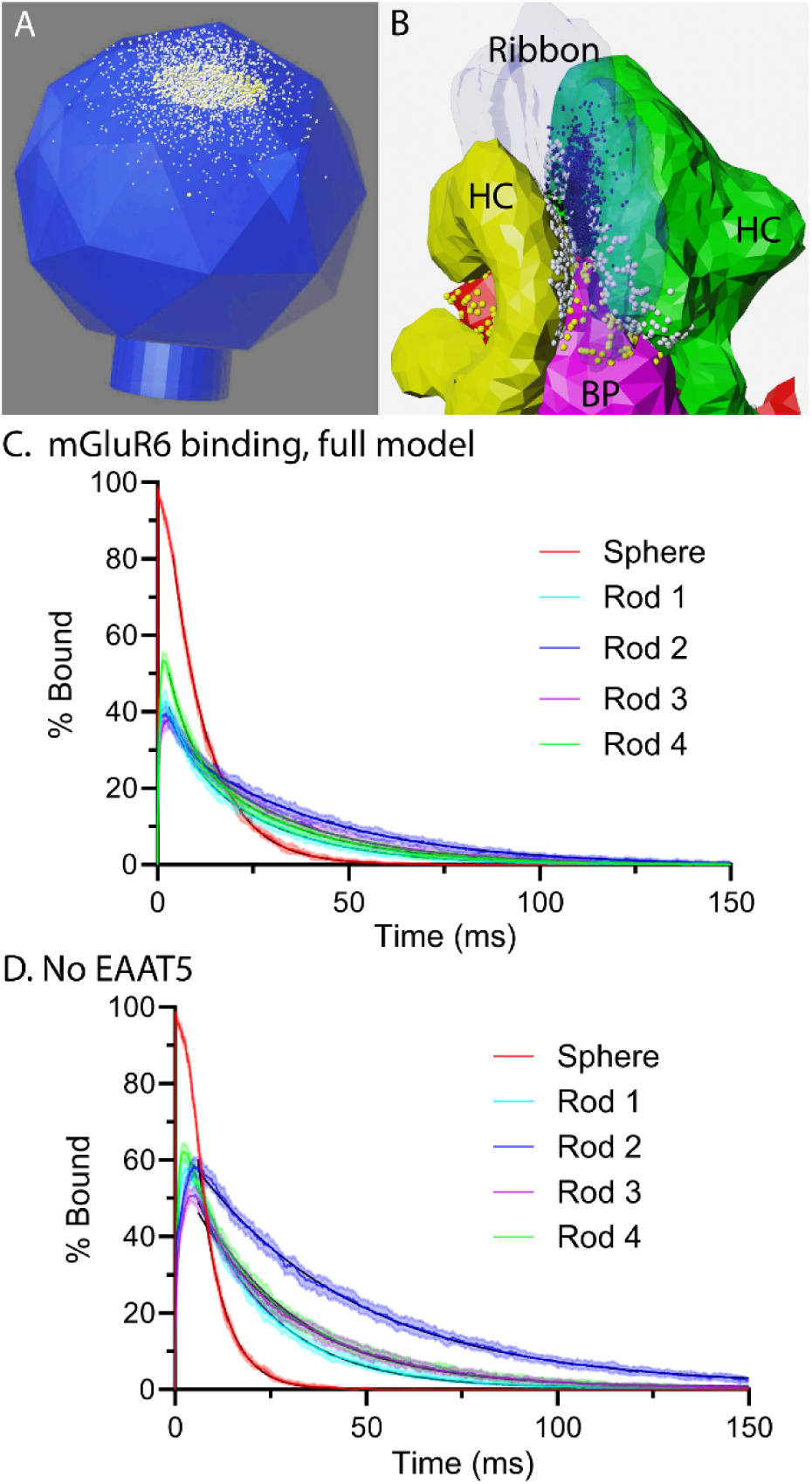
Activation of mGluR6 declines more slowly in rod spherules than a similar size sphere with narrow neck. A. Illustration of the sphere model for rod 2 showing 200 mGluR6 (yellow spheres) in a transparent plane just beneath a release site at the apex. Glutamate molecules following release of a vesicle are shown in white. B. Illustration of rod 1 with mGluR6 (yellow spheres) on bipolar cell (BP) dendrites (purple and red). Glutamate molecules are white when in front of the semi-transparent ribbon and shaded when behind the ribbon. C. Activation of mGluR6 in the sphere and the four rod spherules in the presence of 3,000 EAAT5. The decay time courses in reconstructed rods were fit with two exponentials. Decay in the sphere was fit with a single exponential. Best fit time constants are given in Table 2. D. Activation of mGluR6 in the sphere and four rod spherules without EAAT5. All simulations in this figure are the average of 25 runs using different seed values (± S.D.).

**Table 2.** Best fit parameters from Fig. 6 plotting mGluR6 activity evoked by single vesicle release events in rods 1 to 4, as well as a sphere model.

|  | With 3,000 EAAT5 |  |  |  | No EAAT5 |  |
| --- | --- | --- | --- | --- | --- | --- |
| | Peak % bound | $\tau_{fast}$ (ms) | % fast | $\tau_{slow}$ (ms) | Peak % bound | $\tau$ (ms) |
| Rod 1 | 42.1 | 4.8 | 33 | 24 | 53.9 | 20.4 |
| Rod 2 | 40.6 | 4 | 29 | 40.1 | 58.3 | 42.9 |
| Rod 3 | 39.3 | 7.2 | 18 | 30 | 50.8 | 26.7 |
| Rod 4 | 53.5 | 5.4 | 48 | 28.2 | 56.4 | 26.2 |
| Sphere | 98.6 |  |  | 10.1 | 99 | 6.6 |

We compared these results to the kinetics and peak percentage of mGluR6 activated by a single vesicle in the four reconstructed rod spherules (Fig. 5C). The peak percentage of receptors activated in the two post-synaptic bipolar cells by a single vesicle ranged from 39 to 54%. For these simulations, we averaged the responses of both RBPs together. When we examined each of the two bipolar cells individually, we saw a greater range in the peak level of mGluR6 activation that spanned 19 to 56%, with an average of 47%. In the presence of EAAT5, fitting the decay in mGluR6 activation in rods required two exponentials, with slow time constants ranging from 24 to 40 ms (Fig. 5C, Table 2). In the absence of EAAT5, the decay in mGluR6 activity was well fit with a single exponential. Without EAAT5, mGluR6 activity following release of a single vesicle attained a higher peak value and the response decayed with time constants ranging from 20 to 43 ms (Fig. 5D; Table 2). Rod-to-rod differences and the slower decay of mGluR6 activation in reconstructed spherules compared to a simple sphere illustrate further the influence of synaptic geometry on response amplitude and kinetics.

### Simulations of HC AMPA Receptors

HC dendrites express AMPA receptors (AMPARs) consisting of GluA2 and GluA4 subunits (Hack et al., 2001; Stroh et al., 2018). These two types show similar binding kinetics (Grosskreutz et al., 2003). We simulated HC AMPA receptors using an existing kinetic model for AMPARs (Bartol et al., 2015; Jonas et al., 1993)(Fig. 6A). To assess AMPA receptor kinetics in HCs, we averaged miniature excitatory post-synaptic currents (mEPSCS) recorded from 6 mouse HCs (>50 events per cell). These mEPSCs exhibited rapid rise and decay phases (20-80% rise time = 0.6 ms, τ_decay_ = 0.81 ms; 95% confidence interval: 0.68 to 0.98 ms), as reported previously (Feigenspan and Babai, 2015). We placed 200 AMPA receptors on each of the dendritic tips of the two HCs and simulated the release of a single vesicle containing 2,000 glutamate molecules beneath the ribbon center (Fig. 6B). Simulations in all four reconstructed synapses produced a good match to the actual decay of HC mEPSCs (Fig. 6C). The rise times of simulated mEPSCs were faster than actual mEPSCs, possibly due to imperfect voltage clamp of gap-junctionally coupled HCs (R_s_ = 21.4 ± 5.75 MΩ, R_m_ = 245 ± 201 MΩ, C_m_ = 10.7 ± 8.6 pF, n=16).

**Fig. 6.**
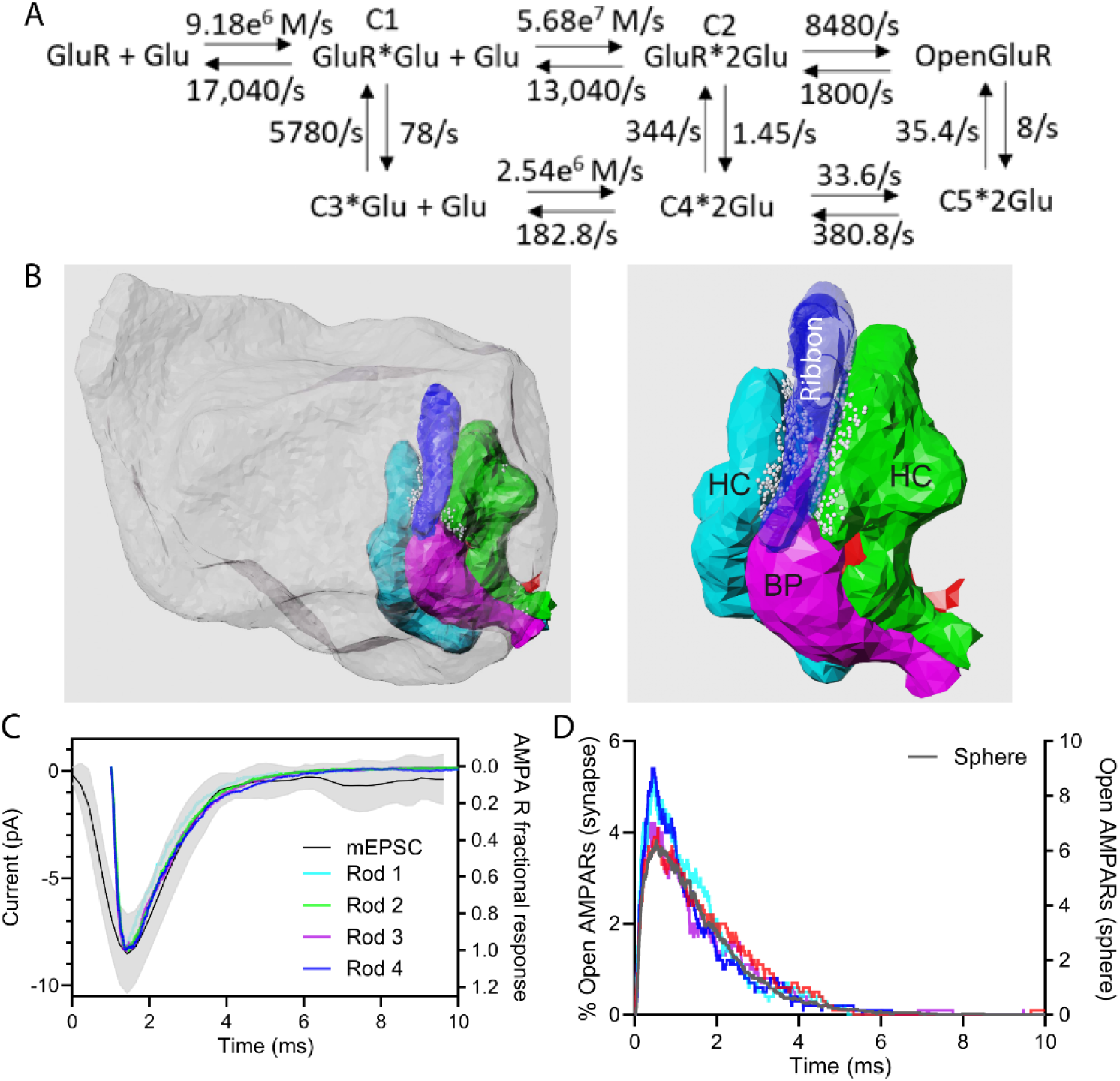
Simulated AMPA receptor activity using a model for GluR2. A. Reaction scheme for AMPA receptor activation from Bartol et al. (2015). B. Illustration of rod 1 spherule with bipolar cells (BP, purple and red), HCs (green and turquoise) and ribbon (semitransparent blue). AMPA receptors are shown as small white spheres. C. Simulated single vesicle AMPA receptor channel openings in four rod spherules compared with the average miniature excitatory post-synaptic current (mEPSC) recorded from mouse HCs (black trace ± S.D.). D. Individual simulations run with 4 different seed values in rod 3 (colored traces). Superimposed on these traces is the average AMPA receptor activity observed with simulations in the corresponding sphere model (n=25 seeds; black trace).

As with mGluR6, we compared simulated AMPAR activation in the reconstructed rod synapses and in the sphere model (Fig. 6D). Fig. 6D illustrates four individual simulations of mGluR6 activity in rod 3 obtained using different seed values. Superimposed on these four colored traces is the average AMPA receptor activity in the corresponding sphere model with 200 AMPA receptors placed in a transparent disc below the apical release site (n=12 seeds; gray trace, Fig. 6D). It is worth noting that unlike mGluR6 where almost half of the receptors are activated by a single vesicle, a much smaller percentage of AMPA receptors is activated. Also, in contrast to mGluR6 activity that showed significantly different kinetics between the sphere and realistic synaptic models, AMPA receptors showed the same kinetics in the sphere and realistic synapse. Synaptic geometry thus has much less impact than intrinsic receptor kinetics on AMPA receptor activity.

### Release site location and dendritic anatomy

Synaptic vesicles can be released at many separate locations along the base of the presynaptic ribbon, suggesting that postsynaptic receptors may encounter widely varying glutamate concentration waveforms depending on their location relative to each released vesicle. We examined the effects of varying release site location on EAAT5 anion channel activity by simulating release at 3 different sites along the ribbon: near the center and at both ends of the length of the ribbon. Within each rod, release at all three sites evoked similar changes in EAAT5 anion channel activity (Fig. 7), suggesting that the observed variability in single vesicle I_A(glu)_ events recorded in individual rods arises primarily from differences in the amount of glutamate released from each vesicle.

**Fig. 7.**
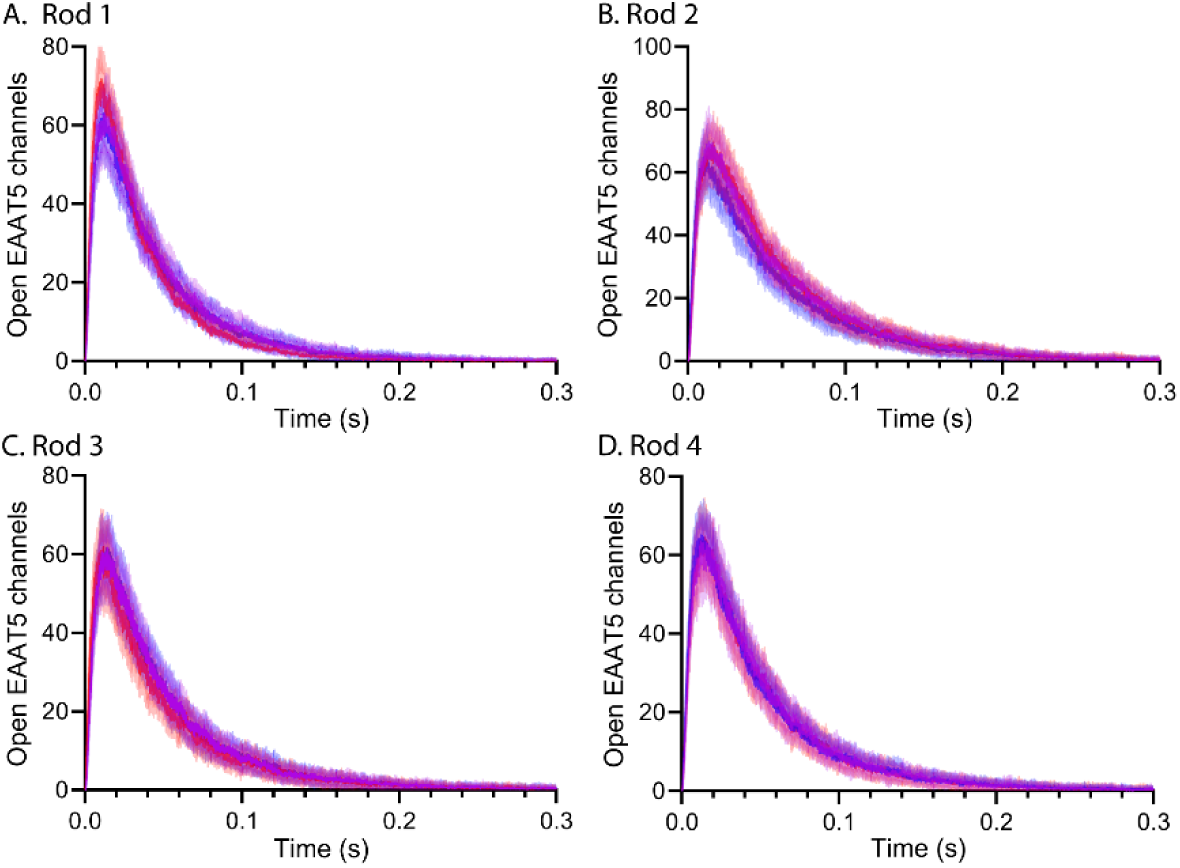
Differences in release site location have an insignificant effect on EAAT5 anion channel activity. In all 4 rods, release was simulated at 3 different sites along each ribbon, one near the center (purple) and at two opposite edges (red and blue). Each trace shows the average ± S.D. of 25 simulations run with different seed values.

We next examined this issue from the standpoint of postsynaptic receptors in RBPs. We examined mGluR6 activity in both post-synaptic RBPs individually and compared six release sites, with three sites on each face of the ribbon (arrows, Fig. 8). Activation of mGluR6 in the two rods varied with release site location. For example, release indicated by the magenta arrow in rod 1 evoked a much smaller response (magenta traces) in bipolar cell 1 (BP1) than release at any other site. Large site-to-site differences remained evident even after we increased α of rod 2 from 0.11 to 0.3 (Fig. 8, insets beneath rod 2). These simulations show that differences in mGluR6 activation arise from geometric tortuosity and are not due to a more tightly confined extracellular space. Some synapses exhibited smaller differences between release sites. For both RBPs contacting rod 4 and BP2 beneath rod 2, mGluR6 activity was similar regardless of release site location. Overall, simulations of single vesicle release events activated 87.4 <u>±</u> 37.1 mGluR6 receptors on each bipolar cell or 43.7 <u>±</u> 18.5% (median: 46.1%) of the 500 receptors placed on each cell (n=48 sites, responses at each site averaged from 25 seeds).

**Fig. 8.**
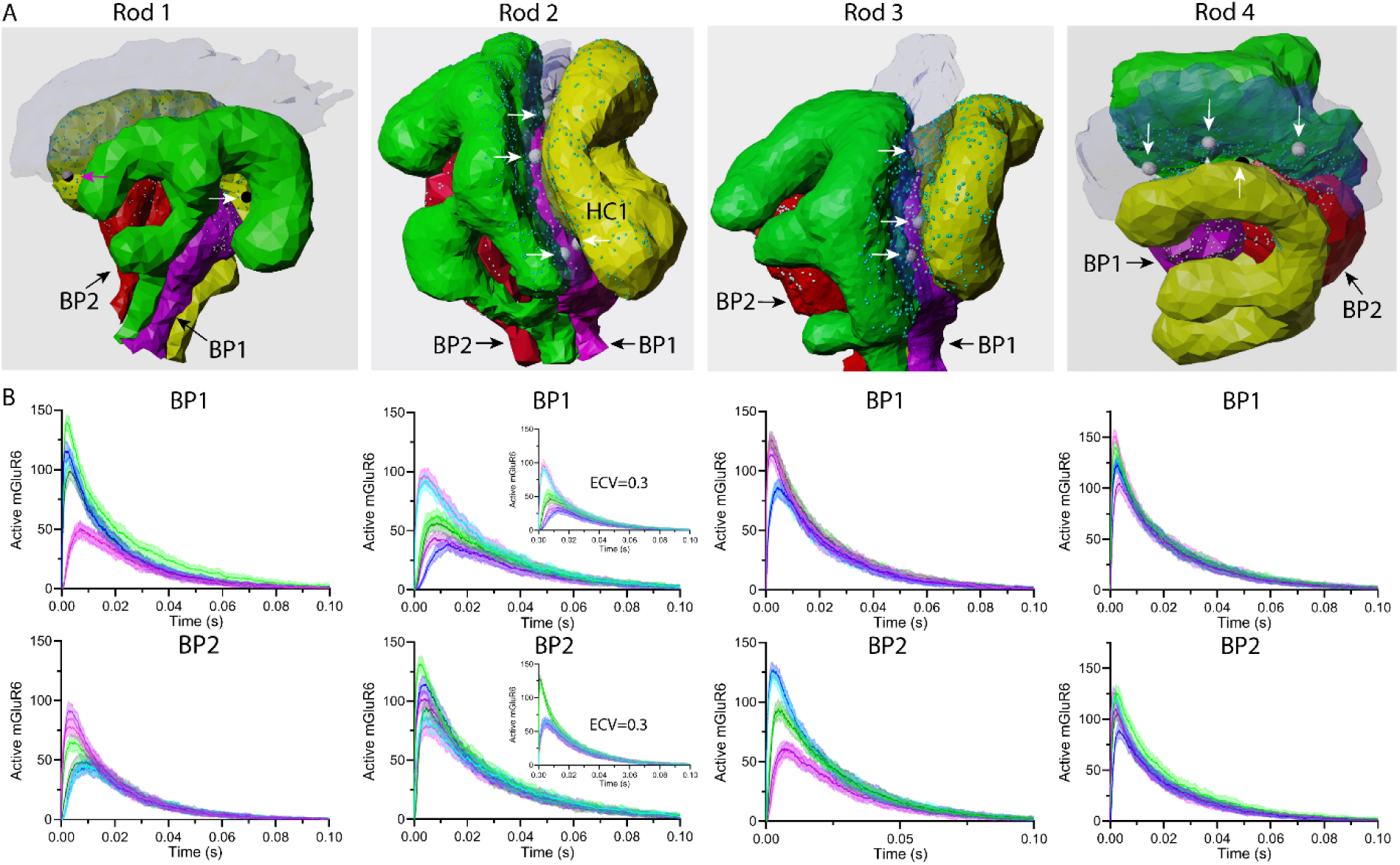
Differences in release site location influence mGluR6 activity. A. Illustrations of rods 1-4 with their post-synaptic cells. Bipolar cells 1 and 2 are shown in magenta (BP1) and red (BP2), respectively. Ribbons are shown in transparent blue. Rod membranes have been removed for easier visualization. Visible release sites are indicated by arrows. The release site corresponding to the magenta trace in BP1 of rod 1 (panel B) is shown with a magenta arrow. B. Plot of the time course of mGluR6 activated by release at six sites, 3 on each face of the ribbon. Responses of the two RPBs beneath each rod terminal (BP1 and BP2) are plotted separately. Insets beneath Rod 2 show the effect on mGluR6 activity of increasing the extracellular volume fraction (ECV) of the synaptic cleft from 0.11 to 0.3. All graphs are the mean ± S.D. of 25 simulations.

AMPA receptor activity in HC dendrites was even more sensitive to release site location (Fig. 9). HCs typically showed stronger responses to release sites placed on the ribbon face nearest to their dendrites. Like bipolar cells, similar site-to-site differences in AMPA receptor activation remained after we increased the volume fraction of rod 2 from 0.11 to 0.3, although the number of active AMPA receptors diminished due to the greater glutamate dilution (Fig. 9B). On average, a single vesicle activated 4.7 <u>±</u> 4.0 % of the receptors (median: 3.7%; n=48 release sites), with a range from 0.14% to 16.5% (Fig. 9F). Fig. 9E shows a histogram of the peak percentage of HC AMPA receptors and RBP mGluR6 activated by release of individual vesicles at different ribbon sites. While single vesicle release events activated a smaller percentage of AMPA receptors than mGluR6, the c.v. for AMPA receptors as a function release site location was larger (0.85) than the c.v. for mGluR6 (0.42).

**Fig. 9.**
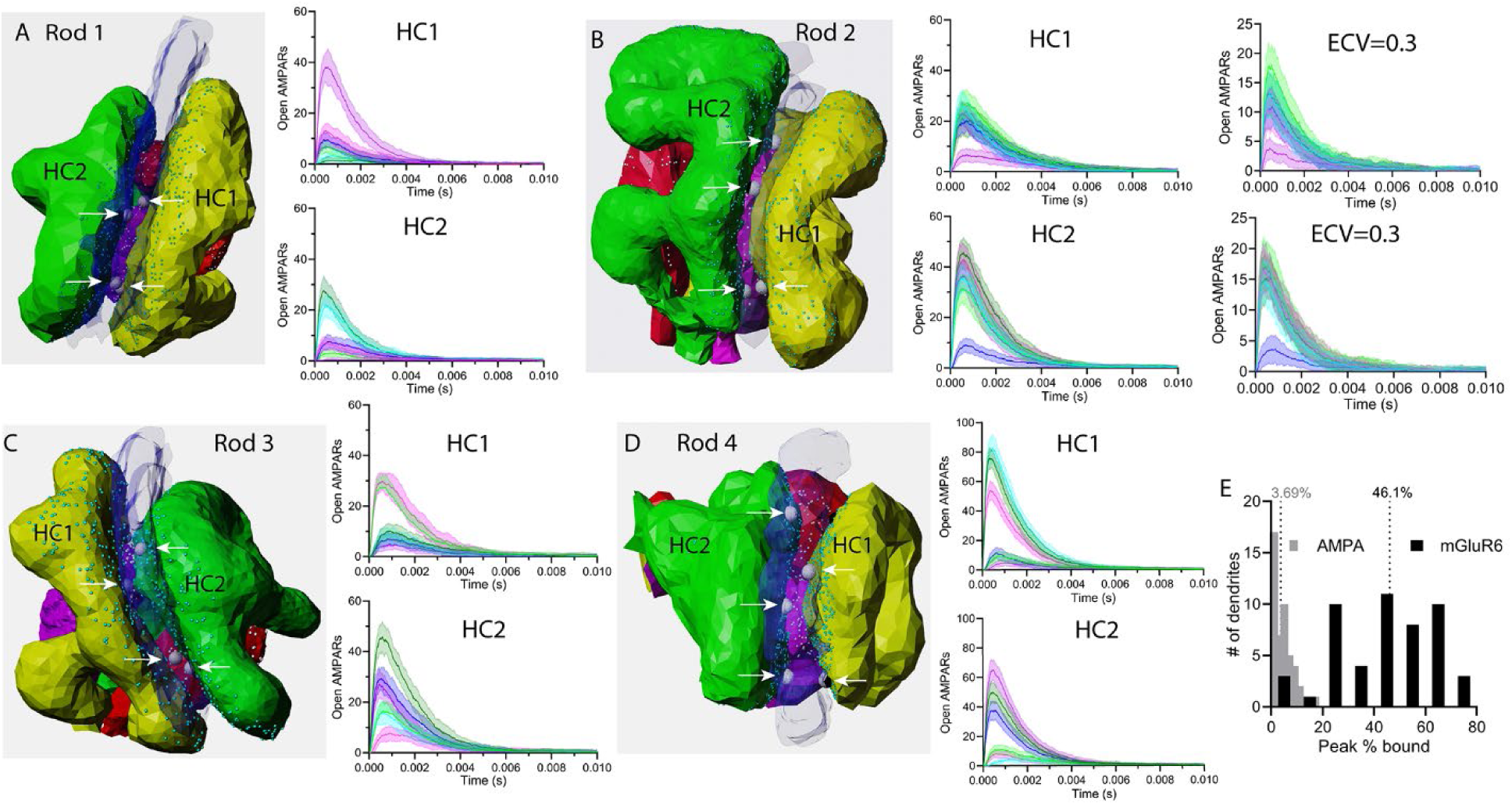
Differences in release site location influence AMPA receptor activity. A. Rod 1 along with its post-synaptic partners. HCs 1 and 2 are shown in yellow (HC1) and green (HC2), respectively. Ribbons are a transparent blue. Rod membranes have been removed for easier visualization. Visible release sites are denoted by arrows. Graphs plot time-dependent activation of AMPA receptors by release at six different sites, 3 on each face of the ribbon. Responses of the two HCs beneath each rod terminal (HC1 and HC2) are plotted separately. B. Illustration of rod 2 with graphs of AMPA receptor activity in HC1 and HC2. Site-to-site variability remained after increasing the extracellular volume fraction (ECV) from 0.11 to 0.3. C. Illustration of rod 3 with graphs of AMPA receptor activity in HC1 and HC2. D. Illustration of rod 4 with graphs of AMPA receptor activity in HC1 and HC2. E. Frequency histogram of the peak percentage of activated HC AMPA (gray) and RBP mGluR6 receptors (black) produced by release at different ribbon sites. For these simulations, we increased the number of AMPA receptors on each HC from 200 to 500. Graphs plot the mean ± S.D. of 25 simulations.

Together, these results show that for glutamate receptors on both RBPs and HCs, release at one ribbon site may have a large effect on one of the two post-synaptic cells and a small effect on the other, while release at a different site may have the opposite pattern. Thus, release site location and dendritic anatomy both have the potential to introduce significant sources of quantal variability at rod synapses.

## Discussion

In this study, we investigated various aspects of synaptic function at the first synapse in the visual pathway using anatomically realistic Monte Carlo simulations of rod spherules. Our results show that the combination of viscous and geometric tortuosity substantially delays glutamate’s escape from the synaptic cleft of the invaginating rod synapse. While glutamate binding and uptake by EAAT5 transporters help to lower glutamate levels, the persistence of glutamate in the invaginating synapse prolongs RBP mGluR6 receptor activity following vesicle release. The simulations also showed the significant impact of differences in cellular architecture and release site location on the amplitude and kinetics of synaptic responses at RBP and HC dendrites.

### Effects of synaptic geometry

For simplicity, Rao-Mirotznik et al. represented the invaginating rod synapse as a sphere with a narrow cylindrical exit (Rao-Mirotznik et al., 1998). Our Monte Carlo simulations of diffusion within that simplified geometry replicated their analytical results showing that glutamate exits such a sphere very rapidly. With smaller mouse rods, glutamate departs a sphere even more quickly, with time constants of less than a millisecond. Replacing the diffusion coefficient for glutamate in saline with a diffusion coefficient that accounts for viscous tortuosity in the extracellular space (Nicholson and Hrabetova, 2017; Nicholson et al., 1979; Nielsen et al., 2004; Rusakov and Kullmann, 1998; Sykova, 2004) slowed diffusion several fold. Replacing the sphere with a realistic synapse incorporating the geometric tortuosity between cells slowed diffusion even further, yielding time constants for passive glutamate diffusion of ~10 ms. This slowing was not due to constriction at the neck or an excessively small extracellular volume fraction in the reconstructed rod spherules. The impact of geometric tortuosity was evinced further by the twofold differences in the rate of glutamate exit among rods with similar cleft dimensions. While most rod spherules are outwardly similar to one another, they can differ significantly in the patterns of their dendritic invaginations (Tsukamoto and Omi, 2022). Our results show that these anatomic differences can have profound effects on glutamate diffusion kinetics.

### mGluR6

To assess the effects of glutamate persistence on RBP responses, we modeled glutamate binding to mGluR6 receptors. Like other class C GPCRs, mGluR6 forms dimers in which agonists must bind both members for full G protein activation (Levitz et al., 2016; Pin and Bettler, 2016). We therefore considered mGluR6 active when bound to two glutamate molecules. Our simulations suggested that release of glutamate from a single vesicle activates nearly half of the mGluR6 receptors on individual RBPs, i.e., at the steepest part of their concentration-response curve, thereby maximizing sensitivity to changes in glutamate release.

Our model did not incorporate downstream signaling pathways engaged by mGluR6 in which glutamate binding triggers the closing of TRPM1 cation channels via interactions involving alpha and beta/gamma G protein subunits (Shen et al., 2012; Xu et al., 2016). Non-linearities in the cascade could also influence response amplitude and kinetics. RBPs employ a non-linear thresholding mechanism in which slight changes in glutamate release cause only slight changes in TRPM1 activation whereas larger changes in release cause disproportionately larger changes in TRPM1 activity (Field and Rieke, 2002; Sampath and Rieke, 2004). This thresholding mechanism filters out small random changes in glutamate release to improve detection of genuine light-evoked changes in release. The threshold for this non-linearity arises from saturation of the intracellular signaling cascade, which was not included in our model. However, our results suggesting that glutamate released from individual vesicles does not saturate mGluR6 are consistent with these previous results.

### AMPA receptors

We found that model parameters developed to fit AMPA receptors in hippocampal pyramidal neurons yielded good fits to decay kinetics of single vesicle mEPSCs in retinal HCs (Jonas et al., 1993). Both types of neurons possess GluA2 receptors (Hack et al., 2001; Stroh et al., 2018). Our simulations predicted a faster rise time than observed in actual mEPSCs, but this could result from limitations in voltage clamp speed of HCs that are strongly coupled to their neighbors.

Unitary mEPSCs in mouse HCs average 3.5 pA in amplitude (Feigenspan and Babai, 2015) and can be generated by opening only 3-5 AMPA receptor channels (Hansen et al., 2021). We achieved a similar number of channel openings per vesicle release event when we placed 200 receptors on each HC dendrite. Comparisons of reconstructed synapses with a sphere model showed that AMPA receptor kinetics are dominated by intrinsic receptor kinetics. The low affinity of AMPA receptors leads to rapid de-activation and this is enhanced by rapid receptor desensitization (Hansen et al., 2021). Kinetic differences among simulated mEPSCs were seen in only a few cases with exceedingly small responses evoked by release at sites distant from individual HC dendrites. Rapid de-activation promotes temporal independence of quanta that promotes linear summation of individual events, contributing to relatively linear contrast-response curves in HCs (Burkhardt et al., 2004). This differs from the steep contrast-response curves in most bipolar cells (Burkhardt et al., 2004). By providing a linear readout of rod and cone membrane voltage, these mechanisms promote linear regulation of photoreceptor output via inhibitory feedback from HCs (Thoreson and Mangel, 2012).

### EAAT5

Studies by Hasegawa et al. (Hasegawa et al., 2006) suggested that presynaptic EAAT5 transporters may retrieve much of the glutamate released by rods. Consistent with a role for this transporter at photoreceptor synapses, genetic elimination of EAAT5 impairs the frequency responses of downstream neurons (Gehlen et al., 2021). Heterologous expression of EAAT5 suggested that uptake by this transporter may be too slow to contribute significantly at rod synapses, but EAAT5 expressed in rods shows fast kinetics suitable for retrieval (Schneider et al., 2014; Thoreson and Chhunchha, 2023). Müller cells whose processes envelope rod terminals (Sarantis and Mobbs, 1992) are also capable of significant glutamate uptake mediated by EAAT1 transporters (Pow et al., 2000; Sarthy et al., 2005). Furthermore, pharmacological inhibition and genetic elimination of EAAT1 impair ERG b-waves that arise from the actions of photoreceptor glutamate release on bipolar cells (Harada et al., 1998; Tse et al., 2014). We used our model to assess the contributions of EAAT5 vs. Müller cell uptake. To do so, we adapted an existing model for EAAT2 to describe the kinetics of EAAT5 anion currents evoked by single vesicle release events in rods (Kolen et al., 2020). Based on the amplitude of evoked and single vesicle I_A(glu)_ events in rods, we concluded that there are ~3,000 EAAT5 at each rod spherule. Following glutamate release, these transporters can rapidly bind up to 3,000 glutamate molecules. Glutamate transport into the rod is slow but nevertheless capable of maintaining glutamate uptake at rates equivalent to 32 vesicle/s. This is close to the peak rate of sustained release by rods of 36 vesicles/s estimated from the size of the readily releasable pool (90 vesicles) and the rate at which that pool can be replenished (2.5/s) (Grabner et al., 2023; Grabner and Moser, 2021; Mesnard et al., 2022a; Mesnard et al., 2022b). While EAAT5 may be able to keep up with much of the release in darkness, glutamate levels nevertheless decline much more rapidly when the synapse remains open and Müller cell uptake is present. The higher glutamate affinity of EAAT1 (2 μM) compared to EAAT5 (10-20 μM) can help to maintain a steep gradient for glutamate diffusion out of the cleft. EAAT2 is concentrated in cones but is also found in rods, where it might contribute to glutamate retrieval (Arriza et al., 1997; Eliasof et al., 1998; Gehlen et al., 2021; Pow and Barnett, 2000; Tang et al., 2022). However, genetic elimination and pharmacological inhibition of EAAT2 have only small effects on dark-adapted ERG b-waves (Harada et al., 1998; Tse et al., 2014) suggesting minor contributions to uptake. Pharmacological inhibition of EAAT2 also has no effect on I_A(glu)_ in rods (Thoreson and Chhunchha, 2023). Our results therefore support the idea that Müller cell uptake retrieves most of the glutamate released at rod spherules but binding of glutamate to EAAT5 helps speed the decline in cleft glutamate levels.

In addition to controlling glutamate levels in the cleft, anion currents activated by glutamate binding to EAAT5 can also modify rod responses directly by altering rod membrane voltage, input resistance, and Ca^2+^ channel activity. The chloride equilibrium potential in rods is ~-20 mV (Thoreson et al., 2003) and so I_A(glu)_ activity in darkness should have a depolarizing effect on rods. However, the stimulatory effects of depolarization are opposed by direct effects of chloride efflux that reduce the open probability of L-type Ca^2+^ channels via actions at specific anion-binding sites (Babai et al., 2010; Rabl et al., 2003; Thoreson et al., 1997, 2000). This inhibitory effect of chloride efflux helps to limit regenerative activation of Ca^2+^ channels and stabilize membrane potential in rods depolarized in darkness. We did not incorporate these and other presynaptic effects into our model.

### Sources of synaptic variability

The earlier conclusion that glutamate diffused rapidly through the invaginating synapse suggested that it equilibrated rapidly throughout the cleft and that post-synaptic dendrites of rod bipolar and HC dendrites all experienced similar glutamate transients in response to individual release events (Rao-Mirotznik et al., 1998). However, our evidence that glutamate exits the invaginating rod synapse more slowly suggests otherwise. Rod-to-rod differences in synaptic architecture led to distinct levels of synaptic activity at both RBPs and HCs. We also observed differences within individual rods in the activity of post-synaptic receptors on RBP and HC dendrites. This variability arises from the fact that, given the same number of glutamate receptors, a more distant RBP or HC will exhibit a smaller response. We also saw differences in response amplitude in individual RBPs or HCs as a function of release site location. Depending on the anatomical arrangement, release at different sites along a ribbon can sometimes produce large differences in mGluR6 or AMPA receptor activity in the same post-synaptic cell. The lower affinity of AMPA receptors made them even more sensitive to differences in release site location than mGluR6. We distributed receptors widely over the tips of RBP and HC dendrites for our simulations. Confining receptor distribution to smaller regions would be expected to produce even more pronounced effects of release site location on receptor activity.

While we saw that mGluR6 and AMPA receptor activity varied with release site location, differences in release site location did not significantly affect the number of EAAT5 anion channel openings. This suggests that much of the variability in single vesicle I_A(glu)_ events in rods arises from variability in the amount of glutamate contained within vesicles. Consistent with this, the range of volumes predicted from vesicle diameters measured in electron micrographs (Fuchs et al., 2014) yields coefficients of variation ranging from 0.28 to 0.44, close to the average coefficient of variation (c.v.) for single vesicle I_A(glu)_ events (0.39) (Thoreson and Chhunchha, 2023). This further implies that much of the variation in the number of molecules per vesicle is explained by variations in diameter, implying that each vesicle contains a similar concentration of glutamate.

### Other invaginating synapses

Many other vertebrate and invertebrate neurons make invaginating synapses, which are particularly abundant in Drosophila (Petralia et al., 2021). However, the invaginating spines and calyceal synapses in other regions of the mammalian brain have a simpler architecture that likely limits effects of geometric tortuosity. At the other end of the spectrum, invaginating synapses of cone photoreceptor cells have an even more complex post-synaptic structure than rods (Sterling and Matthews, 2005) with more than a dozen types of bipolar cells contacting the cone terminal at different sites (Euler et al., 2014). Furthermore, HCs beneath cone terminals have glutamate receptors at both dendrites within the invaginating synapse and on primary dendrites more than 1 micron away from ribbon release sites (Haverkamp et al., 2000). The complex architecture of cone synapses helps to shape response kinetics, with nearby bipolar cell contacts experiencing rapid glutamate changes and distant bipolar cell contacts experiencing slower, smoother fluctuations (DeVries et al., 2006). Along with glutamate receptor properties, the unique architecture of this synapse plays a significant role in the initial filtering and segregation of visual responses into different functionally-specialized, parallel bipolar cell pathways (Grabner et al., 2023).

### Implications for rod release rates and detection of dim light by RBPs

The persistence of glutamate in the invaginating synapse allows greater integration between synaptic vesicle release events. For RBPs, our simulations suggest that mGluR6 receptors may remain active for more than 100 ms after release of a single vesicle. Rao et al. (Rao et al., 1994) suggested that release from rods in darkness must be fast enough so that release events are consistently separated by time intervals shorter than a single vesicle response. Given that single vesicle events last more than 100 ms and assuming Poisson rates of release, this constraint can be achieved with rates of less than 25 quanta/s. However, Rao-Mirotznik et al. (1998) noted that release must also be fast enough to minimize the possibility that a random interval might be mistaken for a genuine slowing of release produced by capture of a single photon. They concluded this required release rates of 100 vesicles/s or more. This exceeds the upper limit on rod release rates of 36 vesicles/s placed by the size of the readily releasable pool and the rate at which that pool can be replenished (Grabner et al., 2023). One possible solution to this apparent dilemma is that release may occur at more regular intervals than predicted by Poisson statistics, thus allowing detection at lower release rates (Schein and Ahmad, 2005). Consistent with this possibility, measurements from rods held at the typical membrane potential in darkness of –40 mV suggest that they release vesicles in multivesicular bursts at regular intervals (Hays et al., 2021; Hays et al., 2020).

Each RBP receives synaptic input from an average of 25 rods (Tsukamoto et al., 2001; Tsukamoto and Omi, 2013). When detecting a single photon event, RBPs must distinguish a small reduction in glutamate release occurring in only one of these 25 rods. Our results suggest several additional sources of synaptic variability that may complicate this already challenging task. These include differences in rod inputs arising from differences in vesicular glutamate content, geometric tortuosity of rod synapses, anatomy of RBP dendrites, location of release sites along the ribbon, location of mGluR6 receptors, and numbers of receptors. How are single photon responses extracted from noise in the face of this variability? In addition to the possibility of regular release, RBPs employ a non-linear thresholding mechanism to extract larger single photon responses from noise (Field and Rieke, 2002; Sampath and Rieke, 2004). It is also possible that retinas will compensate during development for different input strengths by adjusting receptor numbers and/or location to ensure that all the inputs into a RBP are similar. Detailed models of rod spherules offer an opportunity to explore the limitations of these and other potential mechanisms that might be employed for detection of single photon responses by RBPs.

Our simulations revealed surprisingly slow glutamate kinetics at invaginating rod synapses. Slow kinetics allows greater integration of release events at RBP synapses that in turn allows lower release rates to sustain post-synaptic activity in darkness. However, the slow kinetics of glutamate removal introduces additional potential sources of quantal variability by exposing different dendrites to different changes in glutamate. The mechanisms employed by rods to overcome these and other sources of quantal variability and detect light-evoked changes in glutamate release remain to be explored fully.

## Acknowledgments

This work was supported by the NINDS Intramural Research Program (NS003039 to JSD), NIH EY10542 (WBT), EY32396 (WBT), NIH MMBioS P41-GM103712 (TMB), NIH CRCNS R01-MH115556 (TMB), NIH CRCNS R01-MH129066 (TMB), NSF NeuroNex DBI-1707356 (TMB), and NSF NeuroNex DBI-2014862 (TMB). The authors would like to thank Tom Bargar of the Electron Microscopy Core Facility (EMCF) at the University of Nebraska Medical Center for technical assistance. The EMCF is supported by NIH grant 1S10OD026790-01, state funds from the Nebraska Research Initiative (NRI), the University of Nebraska Foundation, and institutionally by the Office of the Vice Chancellor for Research. The authors declare no competing financial interests. This work was completed utilizing the Holland Computing Center of the University of Nebraska, which receives support from the UNL Office of Research and Economic Development, and the Nebraska Research Initiative.

---

**Supplemental Fig. 1.**
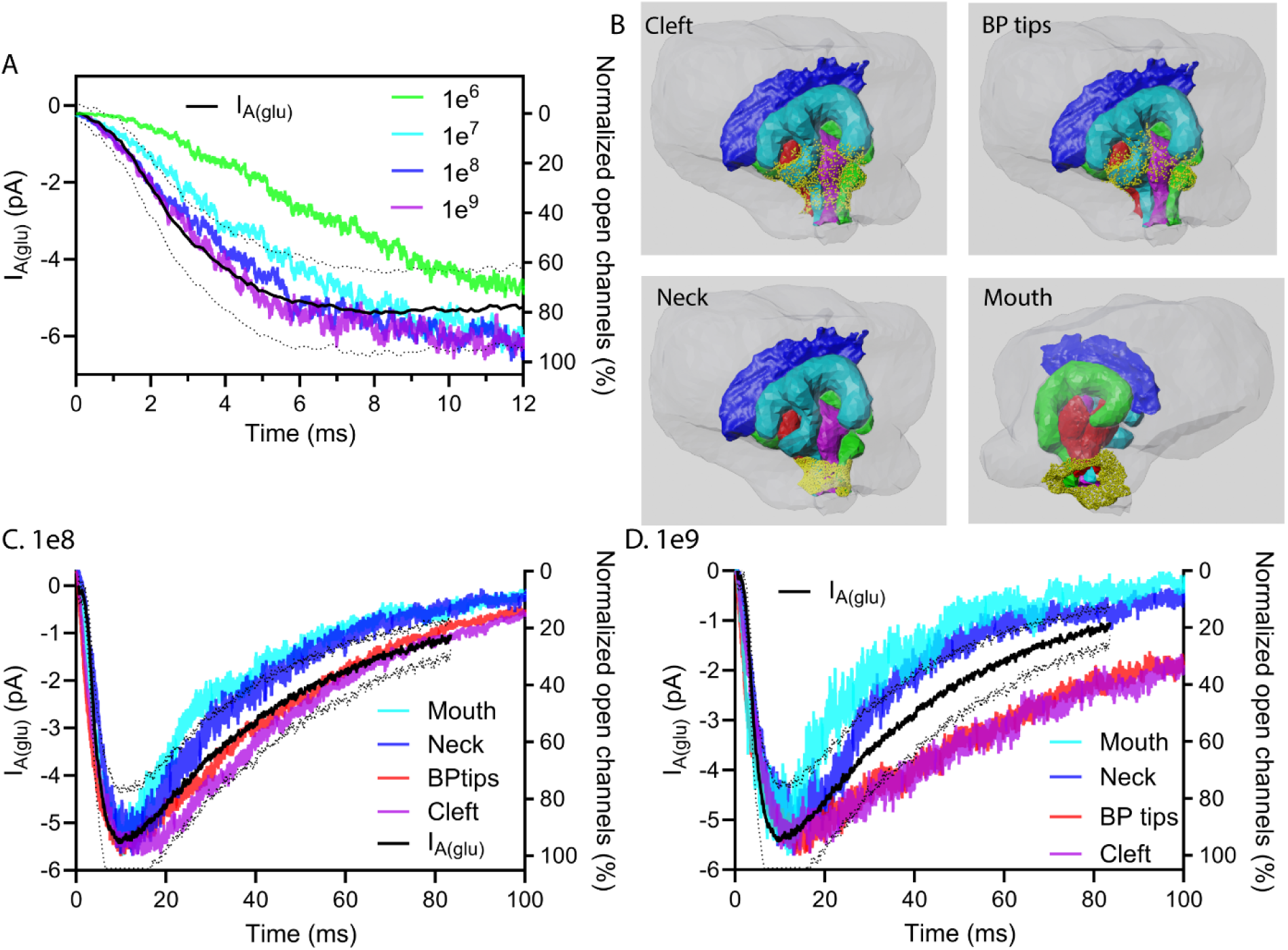
EAAT5 parameters and placement. Comparison of ON rates for glutamate binding to EAAT5 (1 ×10^6^ M/s to 1 × 10^9^ M/s). Black trace shows the average ± S.D. single vesicle I_A(glu)_ in rods. B. Illustration of EAAT5 (yellow puncta) placement in four different regions of the synaptic invagination: throughout the cleft (Cleft), adjacent to bipolar cell dendritic tips (BP tips), the neck of the invagination (Neck), and just outside the mouth of the invaginating synapse (Mouth). Ribbon is colored dark blue. HCs are green and turquoise. RBPs are red and purple. C. Simulated EAAT5 anion channel activity after placing EAAT5 in the different regions shown in B with a glutamate ON-binding rate 1 × 10^8^ M/s. D. Simulated EAAT5 anion channel activity with placement in various locations using an ON rate of 1 × 10^9^ M/S. Traces show the average of 12 simulations run with different seed values.

**Supplemental Fig. 2.**
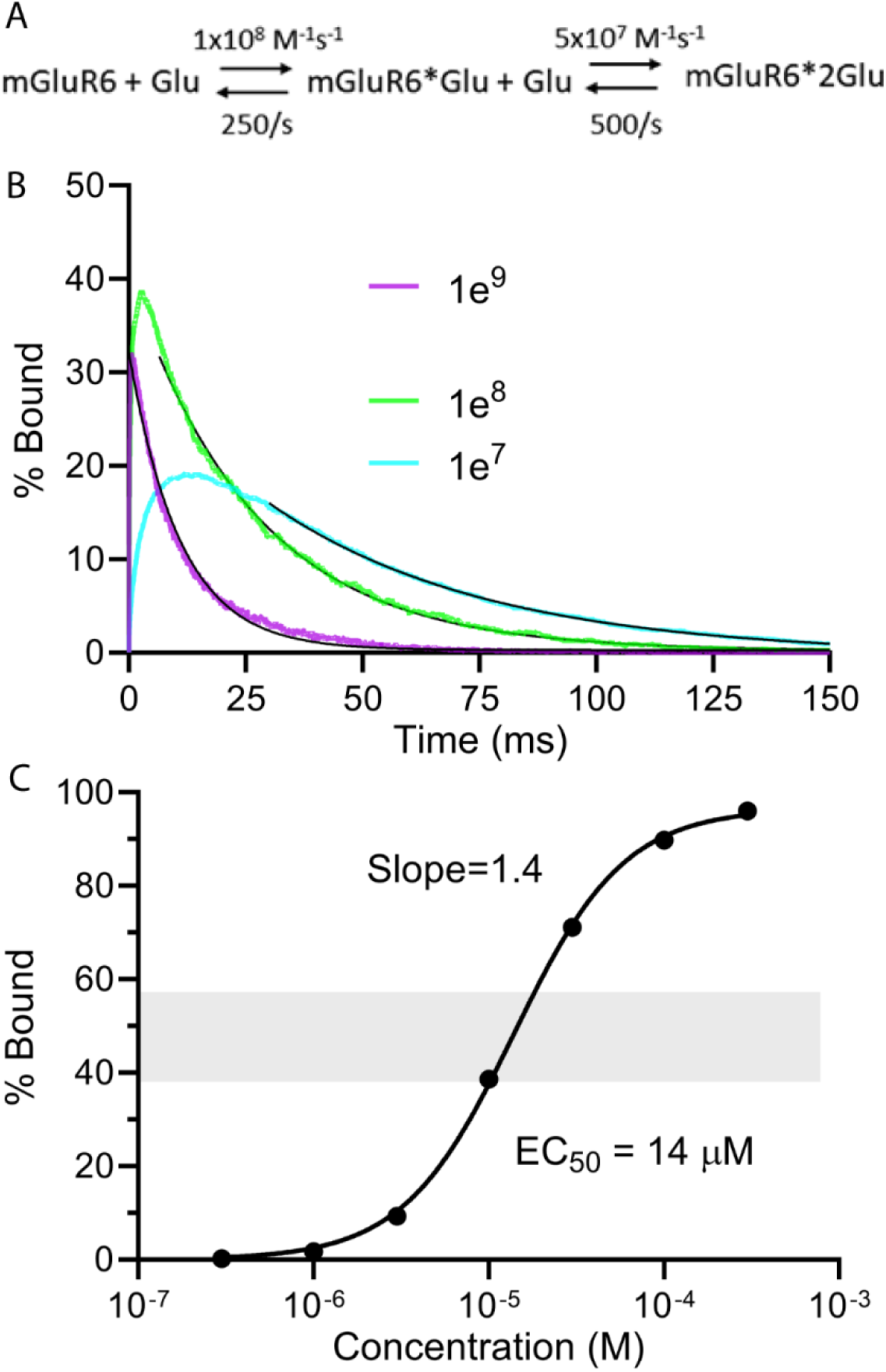
Kinetics of glutamate binding to mGluR6. A. Kinetic scheme for glutamate binding to mGluR6. The activated receptor was considered the doubly bound state (mGluR6*2Glu). B. Kinetics of mGluR6 activation (i.e., doubly bound mGluR6) are shown for different glutamate binding rates. For 1 × 10^9^, 1×10^8^, and 1×10^7^ M/s, the corresponding OFF rates for unbinding of the last glutamate molecule were 2500/s, 250/s, and 25/s, respectively. The later portion of the decay was fit with a single exponential: 1 × 10^9^ M/s, 11.0 ms; 1×10^8^ M/s, 26.7 ms; 1×10^7^ M/s, 46.7 ms. Simulations show the average of 12 seed values run in rod 3 in the presence of 3,000 EAAT5. C. Plot of steady state mGluR6 activation as a function of glutamate concentration using the model parameters in A. Data were fit with a sigmoidal Hill function: EC_50_ = 14 μM. Hill slope = 1.4.

